# Synaptotagmin 7 C2 domains induce membrane curvature stress via electrostatic interactions and the wedge mechanism

**DOI:** 10.1101/2024.01.10.575084

**Authors:** Andrew H. Beaven, Vrishank Bikkumalla, Nara L. Chon, Ariel E. Matthews, Hai Lin, Jefferson D. Knight, Alexander J. Sodt

**Affiliations:** Eunice Kennedy Shriver National Institute of Child Health and Human Development, National Institutes of Health, Bethesda, MD; Postdoctoral Research Associate Program, National Institute of General Medical Sciences, National Institutes of Health, Bethesda, MD; Department of Chemistry, University of Colorado Denver, Denver, CO

## Abstract

Synaptotagmin 7 (Syt-7) is part of the synaptotagmin protein family that regulates exocytotic lipid membrane fusion. Among the family, Syt-7 stands out by its membrane binding strength and stabilization of long-lived membrane fusion pores. Given that Syt-7 vesicles form long-lived fusion pores, we hypothesize that its interactions with the membrane stabilize the specific curvatures, thicknesses, and lipid compositions that support a metastable fusion pore. Using all-atom molecular dynamics simulations and FRET-based assays of Syt-7’s membrane-binding C2 domains (C2A and C2B), we found that Syt-7 C2 domains sequester anionic lipids, are sensitive to cholesterol, thin membranes, and generate lipid membrane curvature by two competing, but related mechanisms. First, Syt-7 forms strong electrostatic contacts with the membrane, generating negative curvature stress. Second, Syt-7’s calcium binding loops embed in the membrane surface, acting as a wedge to thin the membrane and induce positive curvature stress. These curvature mechanisms are linked by the protein insertion depth as well as the resulting protein tilt. Simplified quantitative models of the curvature-generating mechanisms link simulation observables to their membrane-reshaping effectiveness.

## 1. INTRODUCTION

Synaptotagmins (Syts) are a protein family that regulate membrane fusion during exocytosis. Of the 17 mammalian Syt family members, 8 have been shown to trigger membrane fusion in response to Ca^2+^ ions (1, 2). This activity derives from their tandem C-terminal C2 domains, termed C2A and C2B, which each bind two or more Ca^2+^ ions via a cluster of conserved aspartate residues in their calcium binding loops (CBLs). After binding Ca^2+^ ions, the CBLs insert into lipid membranes (3–6). Several models have been proposed for the mechanism by which binding to Ca^2+^ ions and membranes promotes vesicle fusion and exocytosis, but no consensus has been formed (7–17).

Although most Syt mechanistic work has examined the fast neuronal isoform Syt-1 (18–21), this manuscript focuses on Syt-7, which has the strongest Ca^2+^-dependent membrane binding among the Syt family (22, 23). Syt-7’s extreme Ca^2+^ sensitivity derives from a strong membrane affinity of the Ca^2+^-bound state – in particular, Syt-7’s C2A domain binds membranes with much higher affinity and slower release kinetics than its Syt-1 counterpart (22, 24–26). Physiologically, Syt-1 is associated with fast, high-Ca^2+^ release events, such as rapid neurotransmitter secretion, while Syt-7 is associated with slower Ca^2+^ processes, such as asynchronous neurotransmitter release, synaptic facilitation, lysosome fusion, and endocrine secretion of insulin and glucagon (27–34). Syt-7 has also been implicated in transporting cholesterol from lysosomes to peroxisomes (35). In bovine chromaffin cells of the adrenal medulla, Syt-1 and Syt-7 have been observed to exist on separate populations of secretory vesicles, of which the Syt-7 vesicles tend to form exceptionally long-lived fusion pores (36). This observation has led to the suggestion that Syt-7 may stabilize pore structures after fusion in addition to its function in promoting initial fusion upon Ca^2+^ influx (37, 38). Therefore, because of its exceptionally strong membrane affinity and potential for binding stably to fusion pore structures, we have chosen Syt-7 for this initial study of interfacial membrane protein docking, lipid redistribution, and curvature induction on planar membranes.

At a molecular level, Syt-1 (39–42) and Syt-7 (38) strongly associate with anionic lipid membranes, particularly those containing PI(4,5)P2, upon binding Ca^2+^. Furthermore, experiments with Syt-1 have demonstrated that the tandem C2AB (but particularly C2B) disorders *and* demixes phosphatidylserine and PI(4,5)P2 chains (12, 42). Recently, Syt-7 was shown to penetrate more deeply into model bilayers than Syt-1, and that this penetration was critical for its function stabilizing fusion pore open states (43). Given that Syt-7 domains interact with strongly curved membranes during fusion, we seek to understand the interplay of induced membrane disordering (thinning), demixing, and curvature stabilization.

Lipid (dis)ordering is closely related to curvature. In the absence of proteins, lipids that have a propensity to splay their tails outward tend to favor negative curvature (i.e., leaflet curvature is concave relative to the head groups). Therefore, lipid tail splay is an intrinsic trait of negatively curved leaflets. However, curvature-inducing proteins alter both the lipid order and the balance of forces that drive curvature stresses (44–46). One example is the amphipathic alpha helix (AAH), a classic protein element tied to curvature (44). When an AAH is added to a leaflet, the AAH promotes positive curvature (i.e., leaflet curvature is *convex* relative to the head groups) via a wedge mechanism in which the AAH displaces area near the lipid head groups (44–46). This displacement creates a void in the lipid tail region, thinning the leaflet while *stabilizing* lipid splay (44). The correlation of AAH-induced thinning and lipid splay with positive curvature can be understood because the void space underneath the AAH allows adjacent lipid tails space to expand, reducing their negative curvature stress and thus favoring positive curvature. We therefore use leaflet thinning and curvature induction as two signatures of the wedge mechanism (44, 46, 47), even if their observation cannot prove the mechanism is at play. Although Syt-7 is not an AAH, it is known to insert into lipid headgroup regions (48) and could similarly induce positive curvature by creating a void space beneath the embedded protein that disorders and thins the membrane.

Protein-induced local lipid demixing creates unique “lipid fingerprints” that are important contributors to overall membrane association (49, 50). Experiment (51, 52), simulation (53), and theory (54–56) point to the concept that lipid redistribution, and therefore, local membrane material property perturbation, influences fusion pore opening/closure. Particularly in the case of cholesterol, redistribution can cause drastic changes in local pore energetics (53, 56).

Creating fusion pores requires 10s of *k*_B_*T* of energy (57, 58), therefore, proteins are required for fusion pore creation and stabilization on the timescale of regulated exocytosis. However, it is not yet clear how protein interactions might alter the dynamic balance of lipid composition and redistribution that stabilizes the unique curvature profiles at the fusion pore neck. Notably, both leaflets of an ideal fusion pore have net *negative* curvature when measured at the leaflet neutral surface, with the negative principal curvature direction outweighing that of the positive (53, 59, 60). That is, fusion pore necks have negative *Gaussian* curvature, the product of curvature measured in orthogonal directions. Negative Gaussian curvature thins the trans-bilayer hydrocarbon interior (53), similar to the effects of proteins that induce positive curvature via the wedge mechanism. Yet, it is an open question whether membrane-inserting proteins could also stabilize Gaussian curvature, or if the positive curvature induction due to the wedge mechanism will outweigh the thinning effect. Analogously, cholesterol both thickens membranes and favors curvature. We (AHB and AJS) recently found that the thickening preference of cholesterol outweighed its curvature preference, leading to cholesterol depletion at the pore neck (43). Proteins can have a much broader range of membrane interactions compared to cholesterol, and how proteins and lipids work together in the context of a fusion pore is not clear.

Toward this goal, we characterize the membrane interactions of Syt-7 C2 domains that would contribute to a previously proposed (38) “scaffold mechanism” generation (in which proteins form complexes that enforce their curvature onto the membrane) using molecular dynamics simulations of Syt-7 C2 domains on planar lipid membranes. Given fusion pores’ negative Gaussian curvature, we predict that proteins stabilizing the pores will likely have anisotropic curvature preference. Thus, if Syt-7 stabilizes fusion pores through lipid interactions as suggested, it would likely stabilize positive curvature in one direction and negative curvature in another.

Here, we report the orientational distribution, induced local lipid composition, and local membrane deformation of Syt-7 C2 domains, as well as the corresponding curvature stresses induced on planar membranes. First, we examine lipid redistribution around the C2 domains and observe substantial anionic lipid and cholesterol redistribution and contacts with the C2 domains. The C2B domain sequesters more anionic lipid and cholesterol than C2A. Experimental FRET-based assays corroborate the significance of cholesterol, as its presence increases the calcium sensitivity of membrane binding. Further simulations indicate that the C2B domain is embedded deeper in the membrane and lies flatter to the membrane surface than C2A. Finally, we report the leaflet curvature stress caused by the C2 domains as quantified by the lateral pressure profile and find that C2B induces positive curvature, with the (positive-curvature inducing) wedge and (negative-curvature inducing) lipid-attracting mechanisms balancing for C2A.

Taken together, the analysis supports a model in which C2B creates more favorable contacts with the membrane, embeds deeper, disorders lipids more, and promotes stronger positive curvature than C2A. This anisotropic nature of the tandem C2AB domain could be helpful for binding and supporting the anisotropic and varied curvature of a fusion pore. The substantial thinning effect of Syt-7 C2B favors a small fusion pore, perhaps stalling expansion.

## 2. METHODS

### A. Computational system builds and simulations

#### Protein build

The human sequence of Syt-7 was constructed as its components (C2A: residues 134–262 and C2B: residues 266–403) as well as in tandem (C2AB: containing the short link between C2A and C2B). See **Figure 1** for the Syt-7 sequence as well as key residues in the C2A/C2B domains. Independent C2A, C2B, and C2AB were simulated on the planar bilayers.

**Figure 1.**
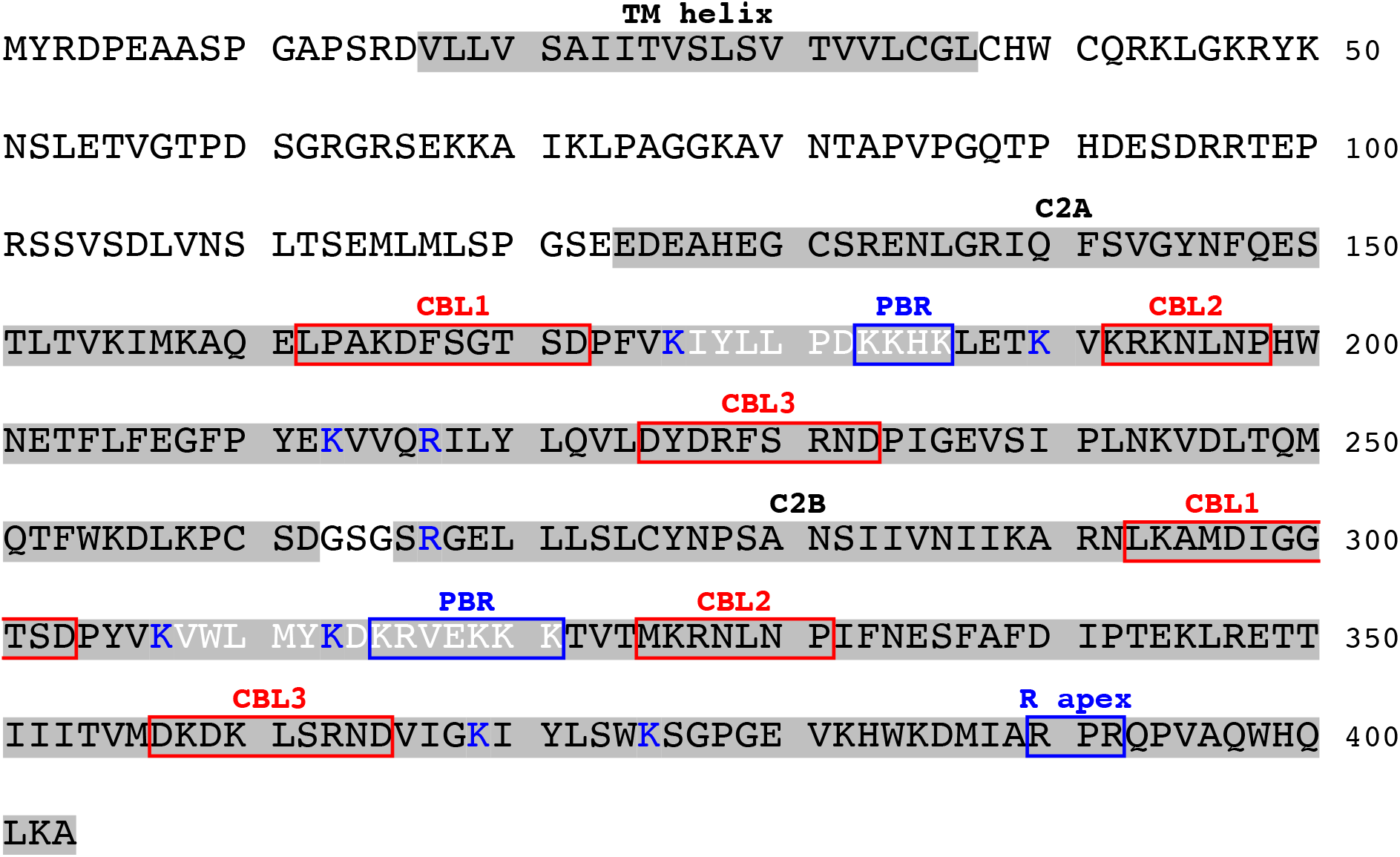
Homo sapiens synaptotagmin-7 sequence from UniProt (ID: O43581) (61). The transmembrane (TM) helix, C2A domain, and C2B domains are accentuated by a grey background. The calcium binding loops (CBLs) of C2A and C2B are boxed in red, and the poly-basic regions (PBRs) are boxed in blue. The arginine apex (R apex) is also boxed in blue. These R apex residues interact with the membrane only in the membrane fusion pore topology (13). Other lysine and arginine residues that form important membrane contacts (see **Figure 8**) are in blue text. The *β*-3/*β*-4 regions of C2A and C2B are in white text (beginning with residues K176 and K307, respectively).

#### Lipid-only and protein-bilayer system builds

Lipid-only and protein-containing planar bilayer systems were constructed using the CHARMM-GUI server and scripts (62). Lipids considered were 1-palmitoyl-2-oleoyl-glycero-3-phosphocholine (PC), 1-palmitoyl-2-oleoyl-glycero-3-phosphoserine (PS), 1-stearoyl-2-arachidonoyl-glycero-3-phosphoinositol (PIP_2_), and cholesterol. These were mixed into bilayers with compositions of PC:PS (75:25 mol%), PC:PIP_2_ (95:5 mol%), PC:PS:PIP_2_ (75:20:5 mol%), and PC:PS:cholesterol (53:17:30 mol%). All systems contained 144 lipids per leaflet, 100 H_2_O per lipid, and 150 mM KCl.

Lipid-only systems were simulated using NAMD (63, 64) with the CHARMM C36m all-atom force field (65, 66) and a 1 fs timestep. PC:PS, PC:PIP_2_, and PC:PS:PIP_2_ systems were simulated for 400 ns apiece, and PC:PS:cholesterol was simulated in triplicate for 165 ns apiece. The temperature was set to 310.15 K by a Langevin dynamics piston (1 ps^−1^ damping coefficient), a semi-isotropic pressure of 1 atm maintained by a Nosé-Hoover Langevin dynamics piston (50 fs period and 25 fs decay time) (67), and covalent bonds involving hydrogen were constrained by SHAKE and SETTLE (68, 69). Nonbonded forces were switched off between 10–12 Å. Long-range electrostatics were calculated by particle mesh Ewald (PME) with a maximum of 1 Å between grid points. These simulations were used to calculate the bending free energy derivative with respect to curvature using the lateral pressure profile (see **Analysis** section).

Protein-containing systems were built with two C2 domains per leaflet (i.e., two C2A, two C2B, or one tandem C2AB) and the lipids described above. Independent C2 domains were initially maximally laterally separated on the bilayer, and they were translated along the *z*-axis to be in minimal contact with the bilayer head groups (**Figure S1**). These systems were briefly equilibrated using NAMD and then converted to Amber format (70, 71) using ParmEd. All C2-membrane systems were simulated in triplicate for ∼2 μs apiece using Amber18’s pmemd.cuda (72–74) to allow for lipid redistribution. The Monte Carlo barostat was set to 1 bar and constant temperature was set to 310.15 K by Langevin dynamics (1 ps^−1^ damping coefficient). Nonbonded forces were switched off between 10–12 Å, long-range electrostatics were calculated by PME, and covalent bonds involving hydrogen were constrained with the SHAKE and SETTLE algorithms. After the ∼2 μs of Amber simulation, the C2-membrane systems were converted back to NAMD and simulated 200 ns with a 1 fs timestep to obtain LPPs. **Table S1** also contains a summary of simulated systems.

### B. Computational analysis

#### Protein definitions

Herein, we used the C2 domain definitions described in MacDougall et al. (38) (also see **Figure 1**). The C2A domain contains residues 134–262 and the C2B domain contains residues 266–403. Within the C2A domain, there are three CBLs: 162–172, 192–198, 225–233. We define the center of mass of the CBLs (CoM_CBL_) by the Ca^2+^-binding aspartic acid residues (166, 172, 225, 227, and 233) of this region. Within the C2B domain, there are also three CBLs: 293–303, 325–331, 357–365 with corresponding Ca^2+^-binding aspartic acid residues (297, 303, 357, 359, or 365) that similarly define the C2B’s CoM_CBL_. Additionally, the C2A domain contains a poly-basic region (PBR) at residues 183–186, while the C2B has a PBR at residues 315–321 and an arginine apex at residues 389 and 391.

Protein height was computed from the CoM_CBL_. Each lipid composition has a different thickness, therefore, to make a balanced comparison across membrane types, the protein location is reported relative to the specific lipid atoms. The C32 atom (carbon bonded to the *sn-2* tail’s carbonyl group) and C3 atom (carbon bonded to the hydroxyl oxygen) were selected for phospholipids and cholesterol, respectively. These two atoms were chosen because: i) C32 is near POPC’s pivotal plane (75); ii) cholesterol’s C3 is a similar height to C32; and iii) C32 and C3 are positioned near the membrane’s hydrophobic surface (see **Figure S2**). As defined, a more positive height indicates a protein bound to the membrane less deeply (i.e., toward the head groups).

#### Protein contact plots

We counted a protein-lipid contact if any residue’s heavy atom (backbone or sidechain) was within 4 Å of a heavy atom of a targeted lipid region (i.e., head group or tail) and/or Ca^2+^. Note, as defined, a residue could be simultaneously interacting with multiple targeted lipid regions of interest at once (e.g., interacting with a PC tail and a PS head group). The interaction was quantified as a “1” if a residue interacted with a targeted region in every frame of the trajectory of the entire ensemble (i.e., four C2 domains per system with each system triplicated). Conversely, an interaction is quantified as a “0” if the residue never interacted with a targeted region.

#### Lipid thickness and distributions

For each simulation frame, for each C2 domain, the entire system was translated so that the CoM_CBL_ was centered at the *xy* origin. A C2 domain’s total CoM (CoM_C2_) was oriented so that the CoM_CBL_ to CoM_C2_ vector lay along the negative *x*-axis. The bilayer’s CoM was centered at *z* = 0. Individual *x*-*y* lipid positions were calculated from the lipid’s CoM. Lipid thicknesses were calculated by the atom definitions described above relative to *z* = 0 (**Figure S2**). In all cases, lipid positions were binned in *x* and *y* with 0.5 Å bins. A bin’s reported thickness is the average of the lipids that resided in the bin. We report the average distribution over the last 1 μs of simulation from the 3 replicas.

#### Free energy derivative with respect to curvature 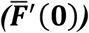

A key component of the analysis is determining the curvature frustration that Syt-7 *induces* into its embedding leaflet. The Helfrich-Canham (HC) Hamiltonian models the energy (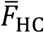; here the bar indicates a *per area* quantity) of leaflet curvature:

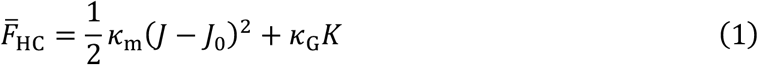

where *k*_m_ is the leaflet bending modulus, *J* is the measured leaflet curvature, and *J*_0_ is the leaflet’s intrinsic curvature. Both *k*_m_ (which sets the energy scale for a deformation) and *J*_0_ are determined by the leaflet’s composition and lateral organization. The constant *k*_G_ is the Gaussian curvature modulus and *K* is the Gaussian curvature. These quantities are important for membrane transitions that require changes in topology (e.g., fusion pore opening).

For planar simulations, information on curvature induction can be obtained by taking the derivative of 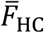 with respect to *J* (the total *J* and *K* of a planar patch are necessarily zero):

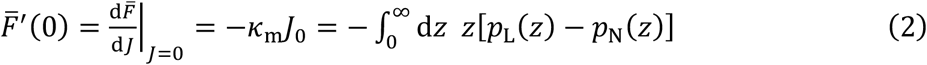

Thus, the derivative is equal to the first moment of the lateral pressure profile (LPP; *p*_L_(*z*) – *p*_N_(*z*)), where *p*_L_(*z*) and *p*_N_(*z*) are the lateral and normal components of the pressure tensor, respectively. The LPP quantifies the pressure experienced at each *z* (with arbitrary graining) through a leaflet (in **Equation 2**, from 0 to ∞). Peaks and troughs in the LPP describe repulsions/attractions in the leaflet, but the ambiguity of the space through which forces act permits only qualitative *local* interpretation of the LPP. However, the zeroth moment of the LPP is the leaflet’s tension, and the LPP’s first moment is equal to the *free energy derivative with respect to curvature* 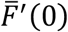; **Equation 2**).

The quantity 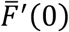 is conceptualized as a “torque” or “leaflet frustration.” It describes the direction that the leaflet would bend if it were unconstrained by hydrophobic and periodic boundary conditions. Because of the negative sign in Equation 2 and naming conventions, a negative 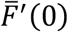 means that a leaflet would bend toward its tails (i.e., a *positive* curvature) and a positive 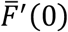 means that a leaflet would bend toward its head groups (i.e., negative curvature). A 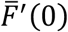 of zero means that the leaflet’s equilibrium average curvature would be planar even if unconstrained by periodic boundary conditions and the opposite leaflet. Curvature caused by a C2 domain is determined by comparing to the lipid-only value (e.g., 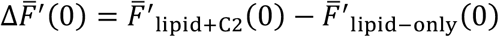). Therefore, a positive 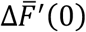 indicates a C2 domain has induced a *negative* leaflet curvature and a negative 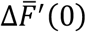 indicates a C2 domain has induced a *positive* leaflet curvature. Alternatively stated, a positive 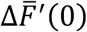 indicates that a C2 domain stabilizes negative curvature, *vice versa*.

### C. Experimental methods

#### Materials

1-palmitoyl-2-oleoyl-sn-glycero-3-phosphoserine (PS), liver phosphatidylinositol (PI), 1-palmitoyl-2-oleoyl-sn-glycero-3-phosphoethanolamine (PE), 1,2-diacyl-sn-glycero-3-phosphocholine (PC), 1,2-dioleoyl-sn-glycero-3-phosphoethanolamine-N-(5-dimethylamino-1-naphthalenesulfonyl) (dPE), cholesterol, and brain sphingomyelin (SM) were from Avanti Polar Lipids (Alabaster, AL). Nitrilotriacetic acid (NTA) was from Alfa Aesar (Ward Hill, MA). All reagents were American Chemical Society grade or higher.

#### Protein and liposome preparation

Syt-7 C2A domain (residues N135–S266), C2B domain (S261–A403), and C2AB domain (N135-A403) were expressed, purified, and dialyzed to remove residual calcium, as described previously (76). Liposomes were also prepared as described previously (76) and were incubated with 10% (v/v) Chelex beads (Bio-Rad, Hercules, CA) overnight at 4 °C to remove residual Ca^2+^. Liposome lipid compositions were designed to approximate the plasma membrane inner leaflet but without PIP_2_, as follows (mol%): with cholesterol, PE:PC:PS:PI:SM:cholesterol:dPE (28:11:21:6:4:25:5); without cholesterol, PE:PC:PS:PI:SM:cholesterol:dPE (28:36:21:6:4:0:5).

### D. Experimental analysis

#### Equilibrium measurement of Ca^2+^-dependent protein-to-membrane FRET

C2 domain liposome binding was assessed using a protein-to-membrane fluorescence resonance energy transfer (FRET) assay in which protein Trp residues serve as the donor and dansyl-modified lipids are the acceptor (77). Buffers were prepared using Chelex-treated Ca^2+^-free water. Quartz cuvettes were rinsed extensively with Ca^2+^-free water before use. Steady-state fluorescence experiments were performed using a Photon Technology International (Birmingham, NJ) QM-2000-6SE fluorescence spectrometer at 25 °C. Excitation slit width was 2 nm; emission slit width was 8 nm. CaCl_2_ was titrated into an initially Ca^2+^ -free solution containing 0.25 μM protein and liposomes (75 mM accessible lipid). Because of the extreme Ca^2+^ sensitivity of Syt-7 C2 domains, a Ca^2+^ buffering system containing 1.5 mM NTA was used for titrations to maintain total calcium concentration ([Ca^2+^]) in excess of protein, as previously described (24). Concentrations of free Ca^2+^ and Mg^2+^ (the latter held constant at 0.5 mM) were calculated using MaxChelator (78). For each titration, FRET was measured (λ_excitation_ = 284 nm, λ_emission_ = 510 nm) over a 10 s integration time for each of three replicate samples. Each intensity value was corrected for dilution, and the intensity of a blank sample containing only buffer and lipid was subtracted. Reversibility was tested by adding excess EDTA after titrations. Normalized data were fitted to the Hill equation,

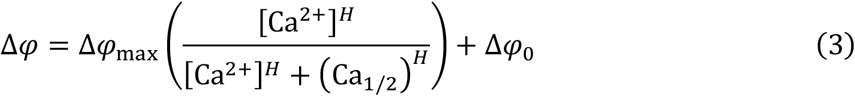

where Δ*φ* is the fluorescence increase, Ca_1/2_ is the [Ca^2+^] at which half of the initially unbound protein becomes membrane bound, *H* is the Hill coefficient, Δ*φ*_0_ is the fluorescence change in the absence of Ca^2+^, and Δ*φ*_max_ is the calculated maximal fluorescence change. Fitting was performed using Kaleidagraph (Synergy Software). Data in figures are shown after normalization of Δ*φ*_max_ to unity for each titration.

## 3. RESULTS

### A. Syt-7 C2 domains sequester PS, PIP_2_, and cholesterol, thin membranes, and disorder lipid tails

Anionic lipids enrich around the CBLs and PBRs/β-4 strands of both C2A and C2B. Enrichment of PS lipids is indicated by red coloring in **Figure 2** (left column) and for both PS and cholesterol of the PC:PS:cholesterol simulations in **Figure 3** (left and center columns, respectively). **Figure 3** shows cholesterol enrichment near the hydrophobic insertion of the CBL, surrounded by a broad region of depletion where PS is enriched. Qualitatively consistent results for PC:PIP_2_ and PC:PS:PIP_2_ are shown in **Figures S3–S4**, respectively. Cross-sectional cuts along the *y* = 0 Å axis of the total leaflet lipid density demonstrate that Syt-7 C2 domains can completely exclude intra-leaflet lipids under the CBLs (**Figures S5–S8**; left). Lipids attempt to fill these voids by tilting (**Figures S5–S8**; right) and thinning/splaying (**Figures 2–3**; right).

**Figure 2.**
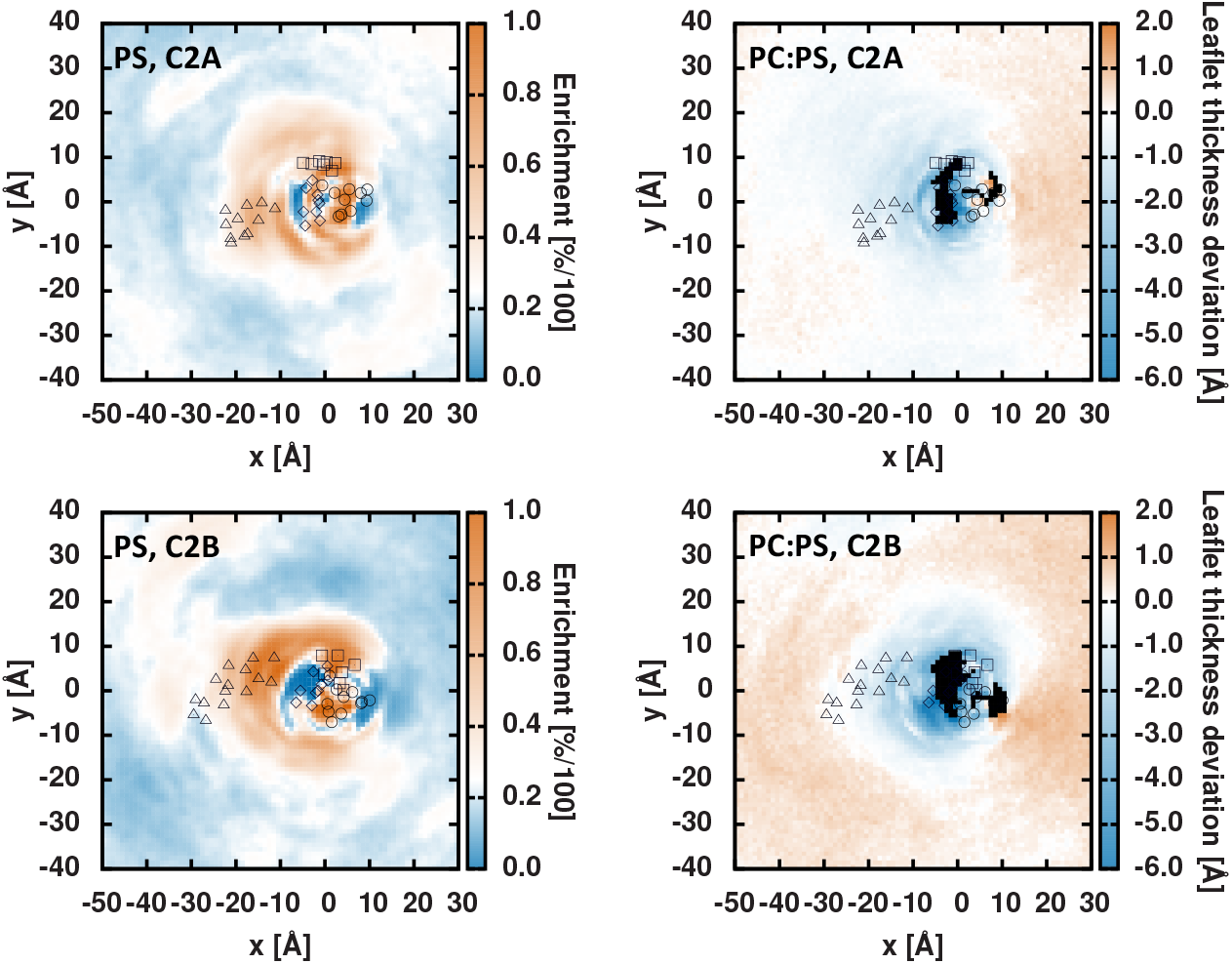
C2A (top) and C2B (bottom) in PC:PS membranes. (Left) PS enrichment relative to the bulk value of 25 mol%. (Right) Leaflet thickness relative to the bulk, lipid-only thickness. For C2A: circles are residues 163–172, triangles are residues 176–186, squares are residues 192–198, and diamonds are residues 225–233. For C2B: circles are residues 293–303, triangles are residues 307–321, squares are residues 325–331, and diamonds are residues 357–365. Black pixels represent bins with total density < 20% of the bulk density.

**Figure 3.**
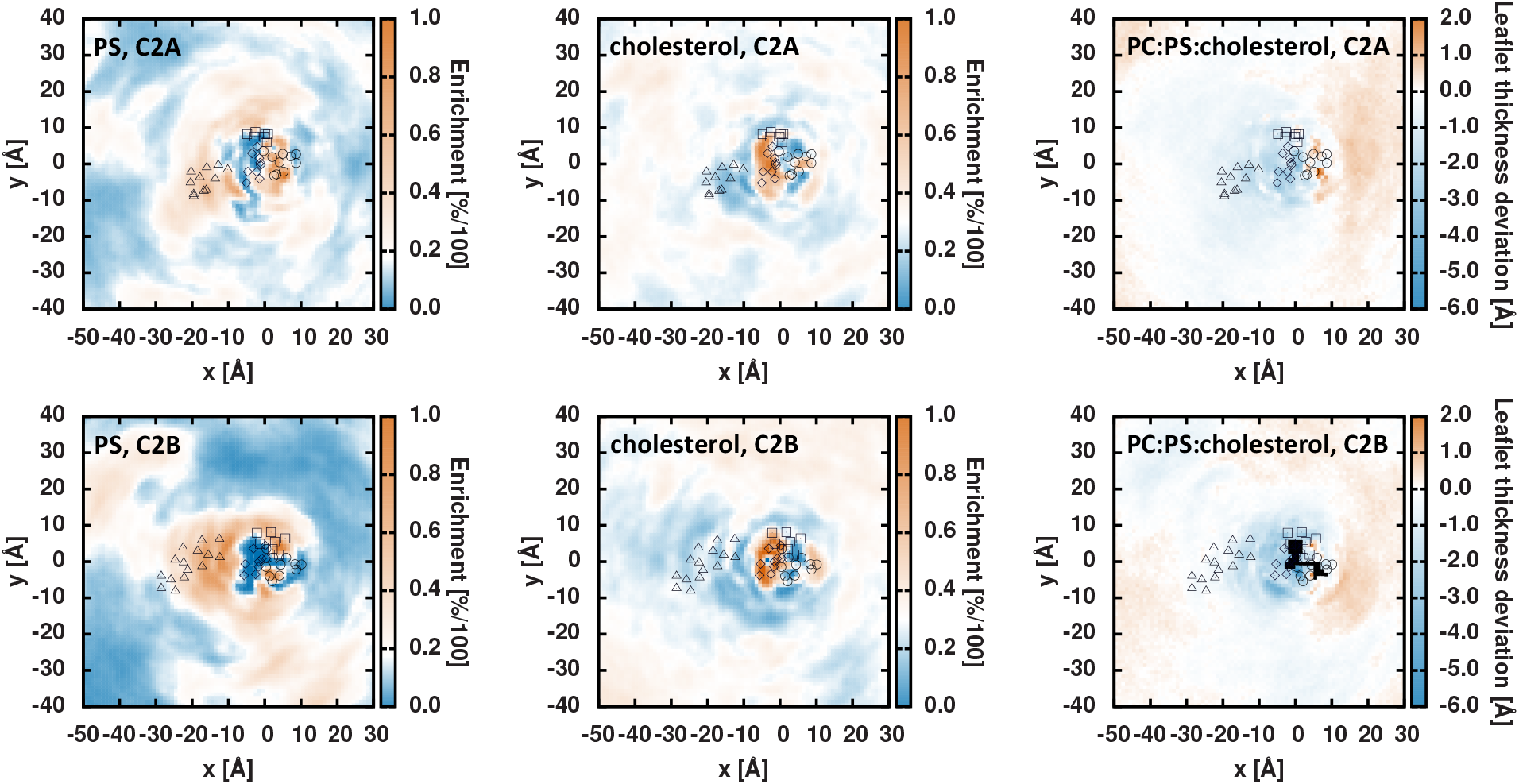
C2A (top) and C2B (bottom) in PC:PS:cholesterol membranes. (Left) PS enrichment relative to the bulk value of ∼17 mol%. (Middle) Chol enrichment relative to the bulk value of ∼30 mol%. (Right) Leaflet thickness relative to the bulk, lipid-only thickness. Black pixels indicate regions where the protein is inserted so deeply that membrane thickness cannot be determined. For C2A: circles are residues 163–172, triangles are residues 176–186, squares are residues 192–198, and diamonds are residues 225–233. For C2B: circles are residues 293–303, triangles are residues 307–321, squares are residues 325–331, and diamonds are residues 357–365. Black pixels represent bins with total density < 20% of the bulk density.

It is well established that Syt-7 C2 domains bind PS (22, 24, 25, 76). However, sequestering cholesterol has not been previously shown for any synaptotagmin C2 domain, to our knowledge. In order to test whether cholesterol enhances membrane affinity for Syt-7 C2 domains, we measured the Ca^2+^ dependence of solo or tandem C2 domains binding to liposomes with or without cholesterol, in a lipid composition that otherwise approximates the plasma membrane inner leaflet without PIP_2_ (76) (**Figure 4**). Removal of cholesterol increased the calcium concentration required to half-saturate liposome binding (Ca_1/2_) by 50% for solo C2A and C2B, and by a factor of two for the C2AB tandem. This magnitude of increase is consistent with a noteworthy secondary effect of cholesterol (relative to PS) on Syt-7 C2 domain membrane binding.

**Figure 4.**
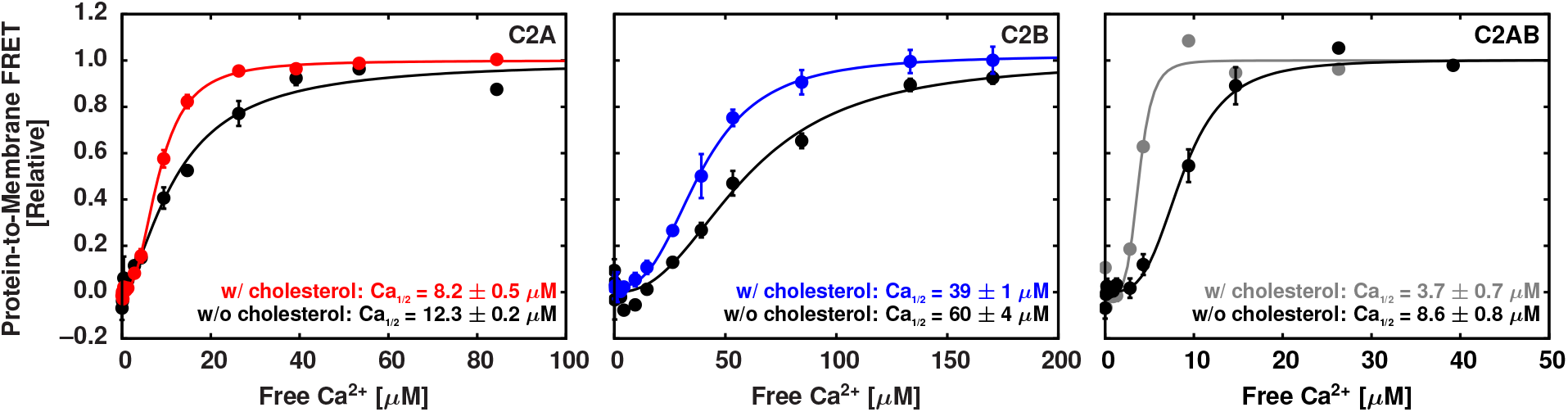
Cholesterol enhances Ca^2+^-dependent liposome binding by Syt-7 C2 domains. CaCl_2_ was titrated into solutions containing the indicated protein domains and liposomes (lipid compositions in **Methods**). Points and error bars shown are mean ± standard deviation of three replicate titrations; where not visible, error bars are smaller than the symbol. Data were fit to Hill curves and the midpoint (Ca_1/2_) values are listed.

### B. Syt-7 C2 domains sequester anionic lipids with the PBRs and wedge with the CBLs

Residue-resolved lipid contacts from simulations also show that both C2A and C2B contact PS and cholesterol. These data are shown in **Figure 5** for PC:PS and for PC:PS:cholesterol in **Figure 6** (PC:PIP_2_ and PC:PS:PIP_2_ data shown in **Figures S9–S10**, respectively). In each contact plot, the C2A solo domain is shown at top left, C2B solo domain at top right, and the tandem domain is shown at bottom.

**Figure 5.**
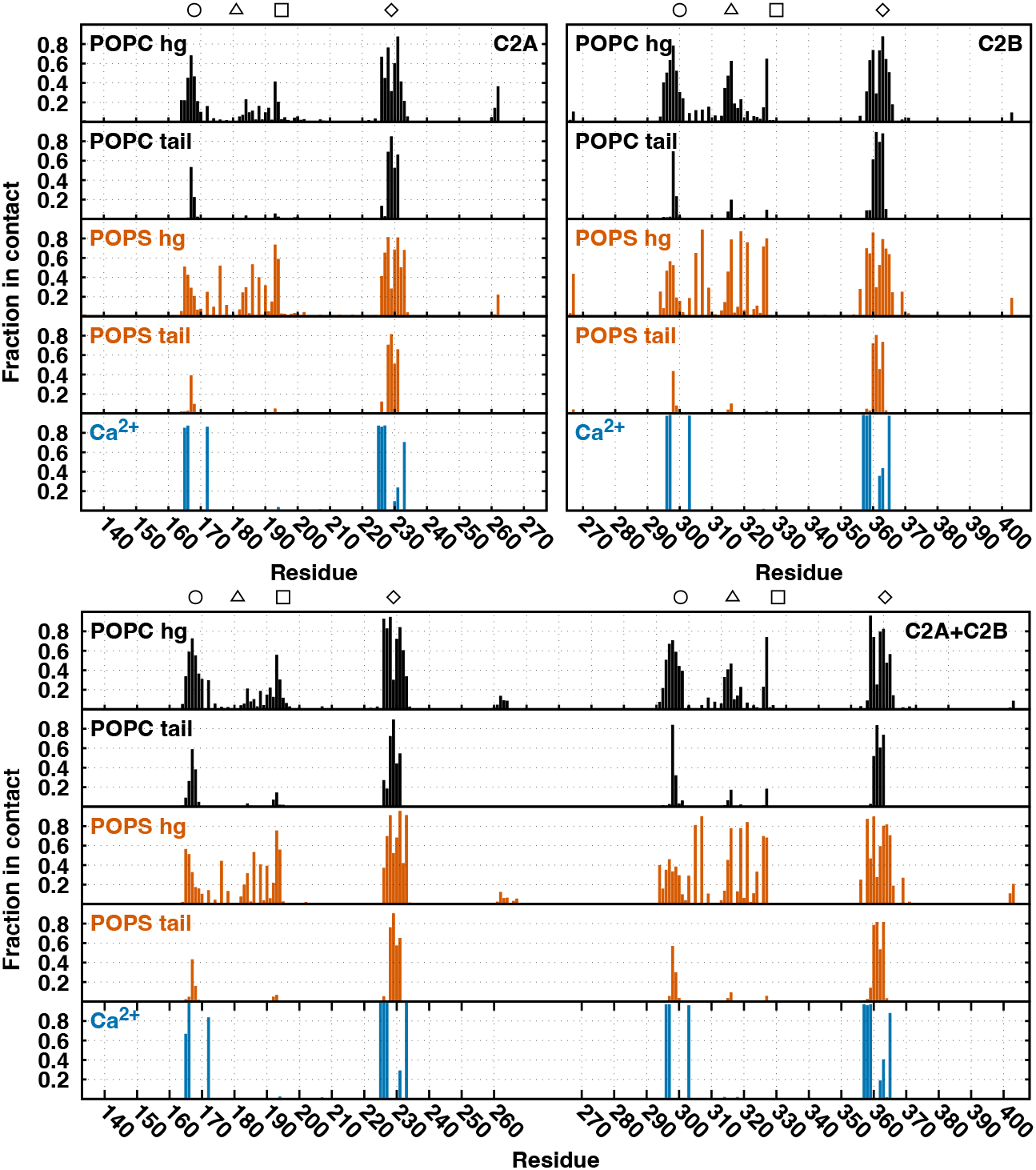
PC:PS contact plots for solo C2A (top left), solo C2B (top right), and tandem C2AB (bottom). PC head group (hg) and tail contacts are in black, PS head group and tail contacts are in vermillion, and calcium contacts are in blue. The circle, triangle, square, and diamond positions correspond to residues 168, 181, 195, and 229 for C2A and residues 298, 314, 328, and 361 for C2B. A value of one indicates that the interaction was present 100% of the time in the ensemble.

**Figure 6.**
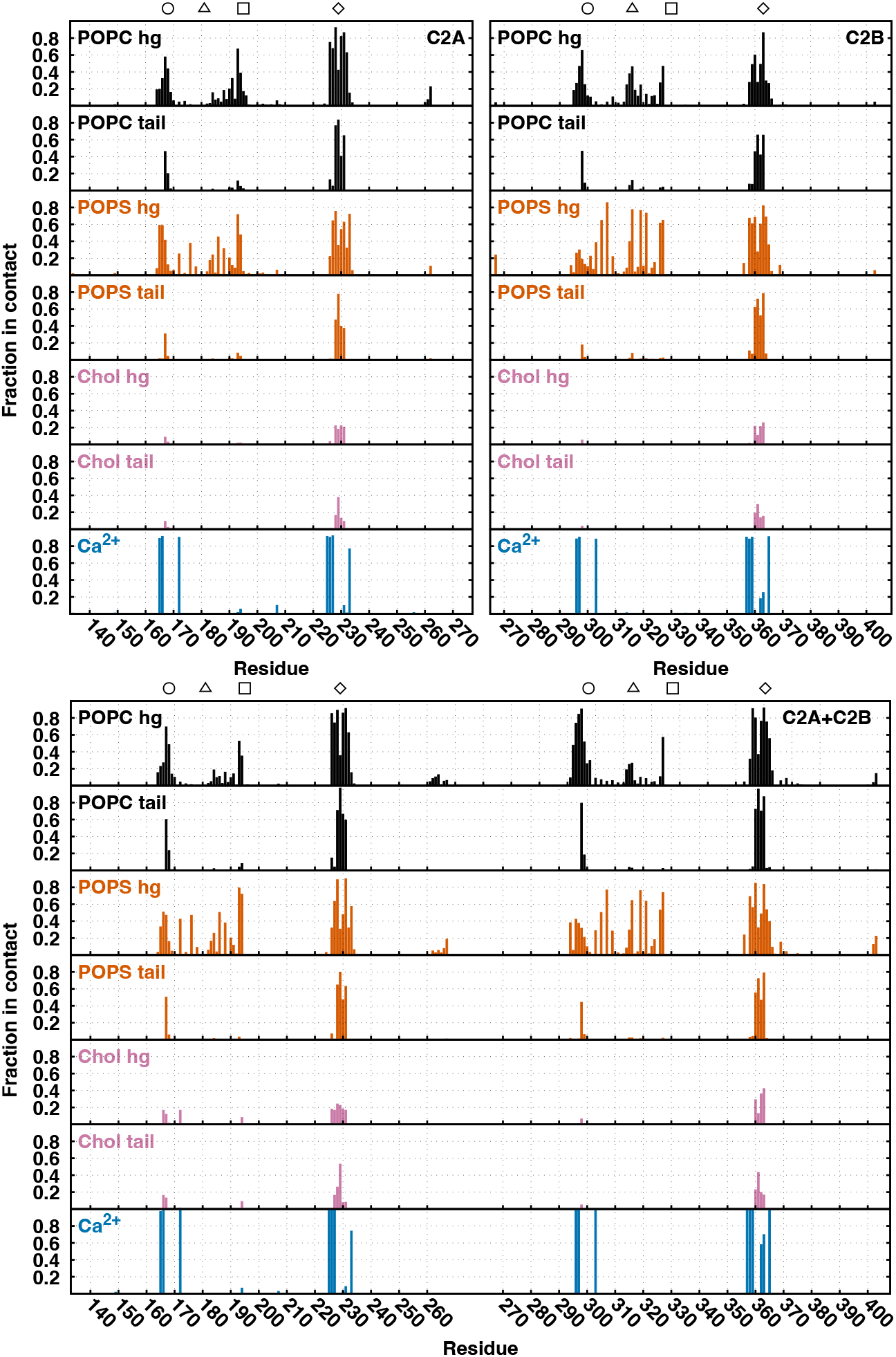
PC:PS:cholesterol contact plots for solo C2A (top left), solo C2B (top right), and tandem C2AB (bottom). PC head group (hg) and tail contacts are in black, PS head group and tail contacts are in vermillion, cholesterol head group and tail contacts are in reddish purple, and calcium contacts are in blue. The circle, triangle, square, and diamond positions correspond to residues 168, 181, 195, and 229 for C2A and residues 298, 314, 328, and 361 for C2B. A value of one indicates that the interaction was present 100% of the time in our entire ensemble.

Cholesterol contacts mostly come from CBL3 (the deepest embedded loop with the most lipid contacts) of both C2 domains. Additionally, there are more total lipid contacts made by C2B’s CBL3 loop in the PC:PS:cholesterol membrane compared to the PC:PS membrane.

**Figures S12–S13** show the cumulatively summed contacts from **Figures 5–6**. Many membrane contacts are formed from the β-3/β-4 strands and the embedded CBLs. Additionally, as expected from the tilt and insertion depth analysis, C2B forms both more contacts with anionic lipids and more total contacts with lipids than C2A. These summed contact results are comparable with those published by Vermaas & Tajkhorshid (Figure 8 of Reference (79)).

As measured by the CBL position relative to the bilayer hydrophobic surface, C2B is inserted deeper (see **Table 1**) into the membrane than solo C2A (consistent with Ref. (79)). Second, we find that the C2B domain in tandem is more deeply inserted than the solo C2B domain in every lipid composition (consistent with Refs. (76, 80, 81)). The C2A domain inserts to approximately the same depth solo or in the tandem domain and is positioned higher than the C2B domain when the two are linked into the C2AB tandem. Inclusion of PIP_2_ makes the C2B domain insert somewhat less deeply, consistent with binding the large PIP_2_ headgroup that sticks up above other headgroups, thus not requiring close approach.

**Table 1.** Protein height (center of mass of the CBLs) in a *leaflet* relative to POPC’s C32 atom. Positive values indicate that the protein is above the C32 atoms. Leaflet thicknesses of 13.45 ± 0.02 Å (PC:PS), 13.24 ± 0.01 Å (PC:PIP_2_), 13.44 ± 0.02 Å (PC:PS:PIP_2_), 16.08 ± 0.01 Å (PC:PS:cholesterol).

|  | C2A (solo) | C2B (solo) | C2A (tandem) | C2B (tandem) |
| --- | --- | --- | --- | --- |
| PC:PS | 10.4 ± 0.3 | 8.5 ± 0.4 | 11.0 ± 0.8 | 7.2 ± 0.3 |
| PC:PIP<sub>2</sub> | 12.4 ± 0.7 | 12.5 ± 0.7 | 11.6 ± 1.0 | 9.0 ± 0.5 |
| PC:PS:PIP<sub>2</sub> | 12.1 ± 0.5 | 10.0 ± 0.5 | 11.0 ± 0.7 | 8.5 ± 0.7 |
| PC:PS:cholesterol | 11.8 ± 0.4 | 10.3 ± 0.4 | 11.4 ± 0.5 | 8.6 ± 0.4 |

### C. C2B tilts to lie flatter on the membrane surface than C2A, while C2A samples a broader tilt distribution

The angle between an internal protein vector (CoM_CBL_ to CoM_C2_) and the *z*-axis quantifies protein tilt (*θ*). A tilt angle of 0° indicates a protein standing straight up relative to the membrane surface (i.e., the protein is aligned with the *z*-axis). A tilt angle of 90° indicates a protein lying flat on the membrane surface. For each system, the first 1 μs of simulation was left off to allow for protein orientational equilibration. Tilts for PC:PS (upper left), PC:PS:cholesterol (upper right), PC:PIP_2_ (lower left), and PC:PS:PIP_2_ (lower right) are shown in **Figure 7**.

**Figure 7.**
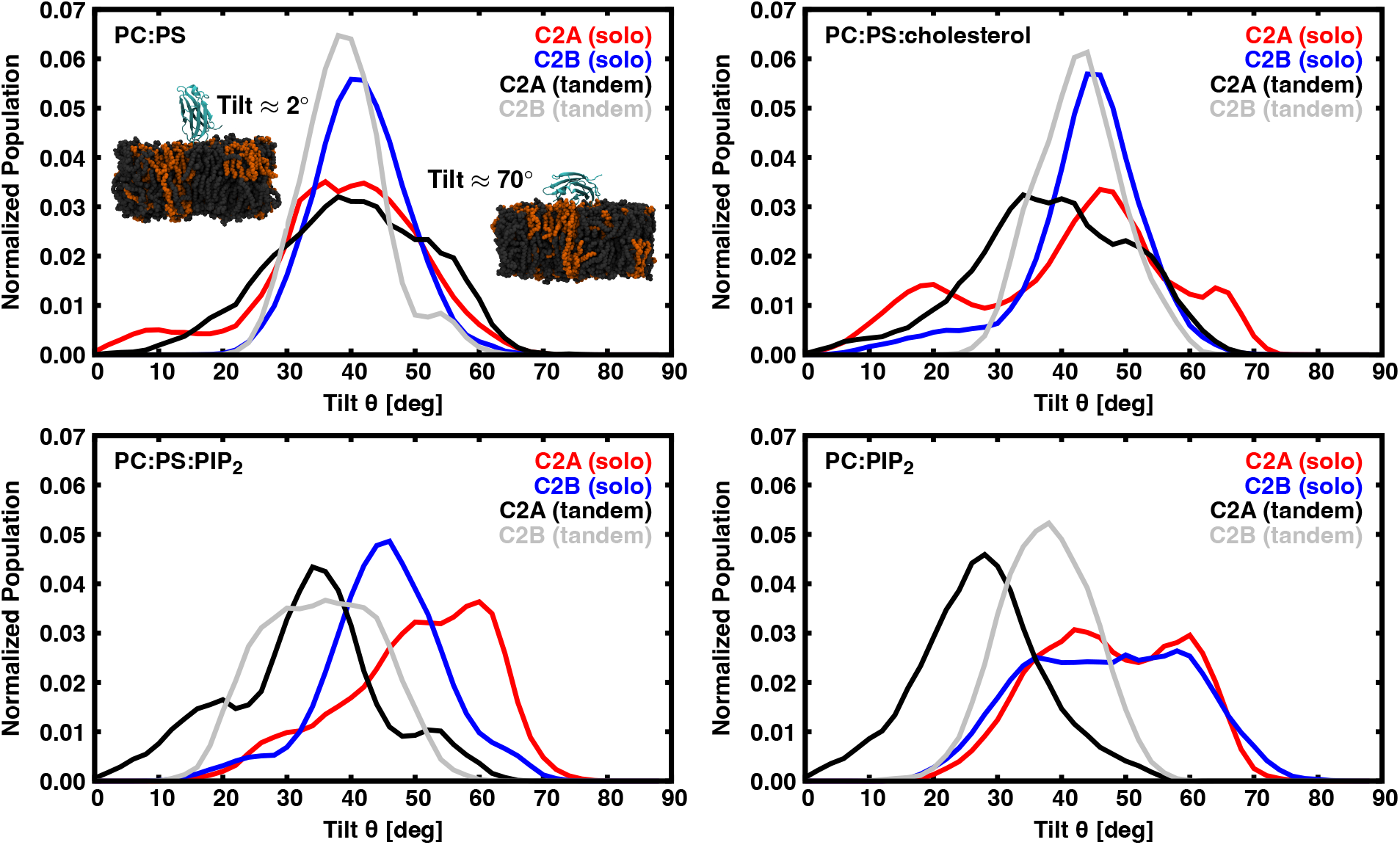
The left and center panels show the individual domain tilts with respect to the *z*-axis when either solo or in tandem (2° bins). Here, a tilt of 0° indicates a domain that is parallel with the *z*-axis (i.e., perpendicular to the membrane surface), while 90° indicates a domain parallel to the membrane surface. On the upper panel inset, molecular images of a C2A domain on a PC:PS membrane (dark grey and vermillion, respectively) show representative conformations. Water and other domains are not shown for clarity.

For systems containing PS, the range of tilt fluctuations for the C2B domain is reduced compared to C2A. It is not immediately clear whether this is structurally related to the varied CBL insertion of the domains, or of the ability of C2B to bind anionic lipids more tightly at its PBR. Being in tandem favors lying flat compared to solo when PIP_2_ is present. This could indicate that the bulky PIP_2_ head groups prevent domains from inserting deeply when solo.

### D. Tilt and lipid contacts are correlated

The correlation between C2 domain tilt (*θ*) and number of contacts a basic residue forms with the membrane (*n*) was calculated and plotted against the residue’s axial position (**Figure 8**) when oriented the same as in **Figures 2–3**. Note that the correlations are not normalized by standard deviation. CBL residues, which are embedded in the membrane, do not strongly affect the tilt. Alternatively, residues far from the CBLs strongly impact the tilt when they form membrane contacts. In **Figure 8**, the steeper slope exhibited by C2A in PC:PS and PC:PS:cholesterol indicates a stronger tilt-contact correlation than C2B. Addition of cholesterol increases the tilt-contact correlation for both C2A and C2B, as evidenced by the steeper slopes in the right panel of **Figure 8**.

**Figure 8.**
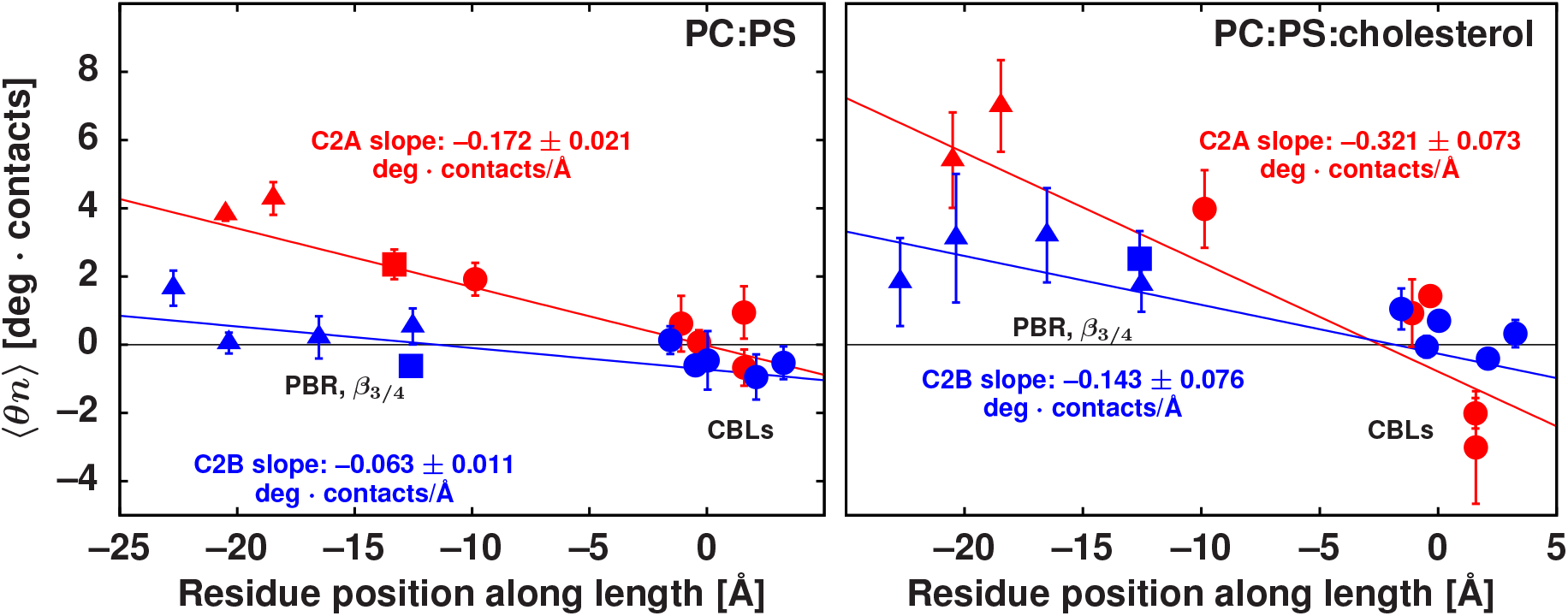
Correlation of tilt and lipid contacts plotted against the axial position along the C2 domain length. C2A is red and C2B are blue. CBLs are circles (C2A = 190, 193, 194, 231, and 228; C2B = 326, 327, 358, 360, and 363). PBRs are triangles (C2A = 184 and 186; C2B = 315, 316, 319, and 321). β-3/β-4 are squares (C2A = 176; C2B= 307). A larger slope indicates a stronger correlation between C2 domain tilt and number of lipid contacts. A correlation of 3 deg · contacts corresponds to an approximate increase of 0.05 contacts per deg tilt, as noted in the Discussion.

### E. Cholesterol modulates Syt-7’s curvature induction on a planar bilayer by relieving positive curvature stress

All 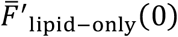 values (without added protein) indicate leaflets with a negative curvature preference. Adding 30 mol% cholesterol (i.e., PC:PS:cholesterol) drastically changes the 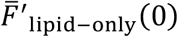 relative to other compositions – demonstrating cholesterol’s strong negative curvature preference in the liquid-disordered phase (82).

Using 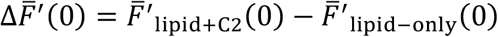, we observe complicated relationships between C2 domain type and membrane composition. Solo C2A induces zero or very weak negative curvature. Solo C2B induces strong positive curvature in PC:PS and PC:PIP_2_, but negligible curvature in PC:PS:PIP_2_ and PC:PS:cholesterol. The curvature induced by the C2AB tandem trends along with the sum of the curvatures induced by the solo domains, that is, it appears to primarily be driven by C2B. Finally, when cholesterol is present, no appreciable curvature is induced by either solo or tandem proteins. We were unable to characterize a statistically significant *dynamic* correlation between 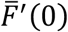 and CBL insertion depth or the number of protein contacts made with the membrane (data not shown).

## 4. DISCUSSION

The above data suggest that Syt-7 C2 domains induce membrane curvature strain by two competing mechanisms. First, the wedge mechanism is driven by membrane insertion and favors positive curvature strain, particularly via the C2B domain. Second, the scaffolding mechanism is driven by protein-lipid contacts distributed over a convex protein surface and favors negative curvature strain as explained below. Differences between C2A and C2B domains in membrane thinning (**Figures 2–3**), insertion depth (**Table 1**), contact surface and orientation (**Figures 5–7**), and tilt-contact coupling (**Figure 8**) suggest that the two mechanisms operate to different extents in these two C2 domains and depend on bilayer composition.

### A. The Syt-7 C2B domain induces positive curvature strain via the wedge mechanism

Syt-7 C2B and C2AB, but not the solo C2A domain, induce positive curvature strain on most of the lipid compositions tested using the lateral pressure profile method (**Table 2**). Consistent with the wedge mechanism, **Figures 2** and **3** (right panels) show a pronounced thinning effect of C2B (bottom) compared to C2A (top) that results from the deeper insertion of C2B. Correspondingly, Syt-7 C2B is more deeply buried than C2A in three of the four lipid compositions for the solo domains, as well as all of the tandem configurations (**Table 1**). Taken together, these results indicate that Syt-7 C2B is an effective membrane wedge that induces positive curvature strain in the absence of cholesterol.

**Table 2.**
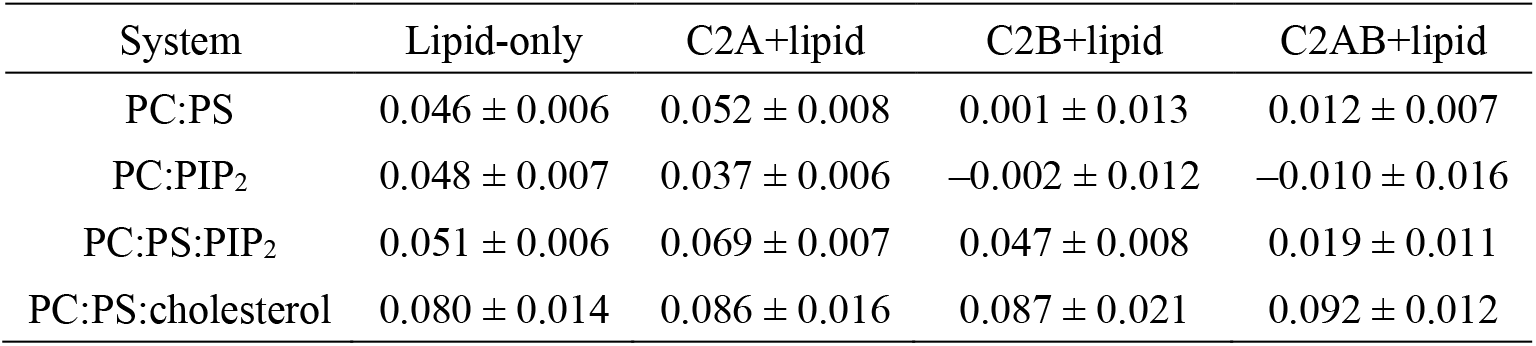
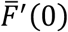 for lipid-only and protein-containing planar bilayers. All units in kcal/mol/Å. Comparison across a row demonstrates Syt-7’s curvature influence into a planar leaflet. A deviation from the lipid-only value indicates curvature induction caused by Syt-7.

| System | Lipid-only | C2A+lipid | C2B+lipid | C2AB+lipid |
| --- | --- | --- | --- | --- |
| PC:PS | $0.046 \pm 0.006$ | $0.052 \pm 0.008$ | $0.001 \pm 0.013$ | $0.012 \pm 0.007$ |
| PC:PIP<sub>2</sub> | $0.048 \pm 0.007$ | $0.037 \pm 0.006$ | $-0.002 \pm 0.012$ | $-0.010 \pm 0.016$ |
| PC:PS:PIP<sub>2</sub> | $0.051 \pm 0.006$ | $0.069 \pm 0.007$ | $0.047 \pm 0.008$ | $0.019 \pm 0.011$ |
| PC:PS:cholesterol | $0.080 \pm 0.014$ | $0.086 \pm 0.016$ | $0.087 \pm 0.021$ | $0.092 \pm 0.012$ |

### B. The deeply inserted Syt-7-C2B forms more lipid contacts than C2A

The regions of the protein surface that interact with lipids are similar in C2A and C2B and include the PBR and three CBLs of each domain (**Figures 5–6 & 8**). However, the C2B domain on average forms a greater number of contacts than C2A in our simulations (**Figures S11–S12**). The increased electrostatic lipid contacts of C2B compared to C2A are also indicated by the enrichment of PS lipids in the vicinity of the PBRs and CBLs (**Figure 2**). The increased contacts are consistent with the C2B domain’s larger disruption of the bilayer (**Figure 2**, right panels). C2B’s deeper insertion favors the CBLs and PBR contacting lipids simultaneously. However, we note that greater lipid contact does not necessarily correlate with stronger membrane affinity. C2B requires a higher calcium concentration than C2A for binding liposomes and was previously shown to have a faster off rate (76) and a weaker affinity (25), at least when PIP_2_ is absent. A possible explanation is that the curvature stress induced by C2B indicates less stable binding.

### C. Syt-7-C2A CBL and PBR lipid contacts are frustrated on a planar bilayer

The Syt-7 C2A domain also inserts into lipid bilayers, albeit less deeply than the C2B domain (**Table 1**). While the C2B domain may insert deeply enough into PC/PS membranes to satisfy lipid contacts in both its CBL and PBR regions simultaneously, the C2A domain does not. Thus, the C2A domain samples a wider range of tilt angles than C2B (**Figure 7**). The strong correlation between C2A domain tilt and lipid contacts suggests that there is competition between forming contacts with its CBL region or its PBR region. Given that the tilt/contact correlation is greater for C2A than C2B (**Figure 8**) and that C2A has fewer lipid contacts on average (**Figures 3, 4**, and **5**), the balance for C2A appears to favor CBL contacts more than for C2B, giving rise to a population of C2A domains with low tilt angle (**Figure 7**). This low-tilt state may be stabilized by the greater hydrophobic character of the phenylalanine residues on the CBLs of C2A compared to the isoleucine/valine residues of C2B, and/or by the fact that the C2A domain’s PBR is less electropositive than that of C2B.

### D. Cholesterol impairs deep insertion and wedging

The presence of cholesterol weakens the membrane thinning effects of both C2A and C2B domains (compare **Figure 2** to **Figure 1**) and correlates with shallower membrane insertion (**Table 1**), consistent with the known membrane condensing effect of cholesterol (83). Addition of cholesterol in our simulations also shifts the insertion and contact profile of the C2B domain closer to that of C2A. The solo C2B domain samples more low-tilt orientations in the presence of cholesterol (**Figure 7**) and its tilt-contact correlation becomes significant (**Figure 7**). Our experimental measurements (**Figure 4**) show that both domains bind liposomes better (more Ca^2+^-sensitive) in the presence of cholesterol. As our simulations indicate that cholesterol is enriched directly beneath the CBL regions, we hypothesize that cholesterol enhances CBL membrane affinity and thereby increases the binding frustration between the CBL and PBR regions to some extent for both C2A and C2B domains. In this interpretation, insertion of the CBL induces stress in the bilayer that is reduced by the enriched cholesterol, which fills the void created underneath the CBL.

### E. A tilt-contact model of Syt-7 scaffolding predicts C2 domain curvature induction

Negative curvature would reduce the conflict between PBR and CBL lipid contacts by completing potential protein-lipid contacts. This effect is reflected in the derivative of the free energy with respect to curvature. The observation that C2A does not induce significant negative curvature as measured using the LPP method (**Table 2**) may be a result of competing wedge and tilt mechanisms; the C2A domain also inserts (**Table 1**) and thins membranes (**Figure 1**) consistent with the wedge mechanism. Nevertheless, the curvature induction of C2A on PC:PS is significantly more negative than that of C2B, for which correlation of tilt and lipid contacts is much smaller. Inclusion of cholesterol lessens the insertion depth (**Table 1**) and positive curvature stress induction (**Table 2**) for C2B and increases its tilt-contact coupling (**Figure 8**). The tilt-contact correlation effect is discussed below where a simple model is described for how Syt-7 C2 domains couple to fusion pore curvature.

#### Curvature induction by PBR-lipid contacts

To model the influence of Syt-7 C2 domains on fusion pore energetics requires quantifying the curvature-generating force. The simulations here provide two complementary pieces of information. First are the *molecular* details about the interactions of the domains (thinning near the insertion of the C2 domain, the variation of lipid-PBR contacts with tilt). Second is a quantification of the force itself by means of the LPP, which provides the derivative of the free energy with respect to curvature. To predict how changes in structural features lead to changes in the force requires connecting the structural features to the quantitative force through a logical model.

Consider first the coupling of C2 domain tilt and PBR-anionic lipid contacts. This analysis assumes that tilting increases contacts (**Figure 8**) by decreasing the distance of the PBR to the leaflet surface. *Curvature* acts equivalently, bringing the leaflet closer to the PBR domain at constant tilt (see the left panel of **Figure 9**, where the grey leaflet curves to bring anionic lipids close to the PBR). The distance *z* between the surface and PBR as a function of protein tilt *θ* goes as:

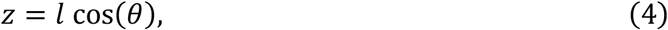

where *l* is the distance of the PBR residue from the pivot of tilting (ca. 15 Å, see **Figure 8**). Curvature (*J*) similarly relates the height of the anionic lipids as a function of *l*:

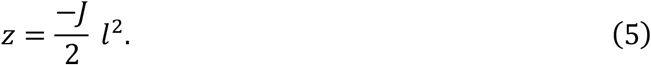

A schematic illustrating **Eqs. 4** and **5** is provided in **Figure 10**. The number of contacts per degree theta (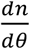 **Figure 8**) is then used to estimate the number of additional contacts per unit curvature under the assumption that contacts are determined only by *z*, and thus that *θ* can be represented as a function of *z* (concerning the computation of the correlation of contacts and *θ*):

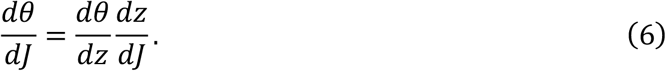

For normally distributed *θ* and *n*, the derivative 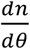 can be estimated from the correlation ⟨*n θ*⟩ as 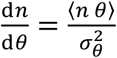, where 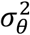is the variance of *θ*. The variance of *θ* for C2B in PC+PS (the distribution most closely resembling a normal distribution, see Fig. 7, top left) is 54 deg^2^. The free energy change with curvature can then be estimated as

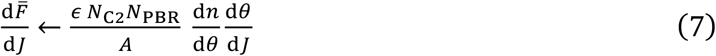

where *ϵ* is the energy per contact, *N*_PBR_ is the number of positively charged residues in the PBR, *N*_C2_ is the number of C2 domains per leaflet in the simulation, and *A* is the area of the box (normalizing the free energy per unit area).

Note here that to compare to the simulation values in Table 2, the free energy is normalized per unit area. For C2A, simulated with *N*_C2_ = 2, and using *N*_PBR_ = 4, contacts with an average 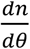 of 0.05 contacts per degree (see **Figure 8**) yield an expected change in the first moment of the lateral pressure profile of 0.03 *ϵ*, corresponding to the difference between C2B and C2A if the strength of a lipid contact is roughly 1 kcal/mol. Wider variance in the C2A angle distributions may weaken 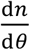 somewhat, increasing the lipid contact strength necessary to explain the free energy derivative. Due to the undetermined contact energy the model cannot be made perfectly quantitative from first principles. Yet the calculation indicates that a reasonable interaction energy is sufficient to explain the relative negative curvature induction of the C2A domain over C2B.

**Figure 9.**
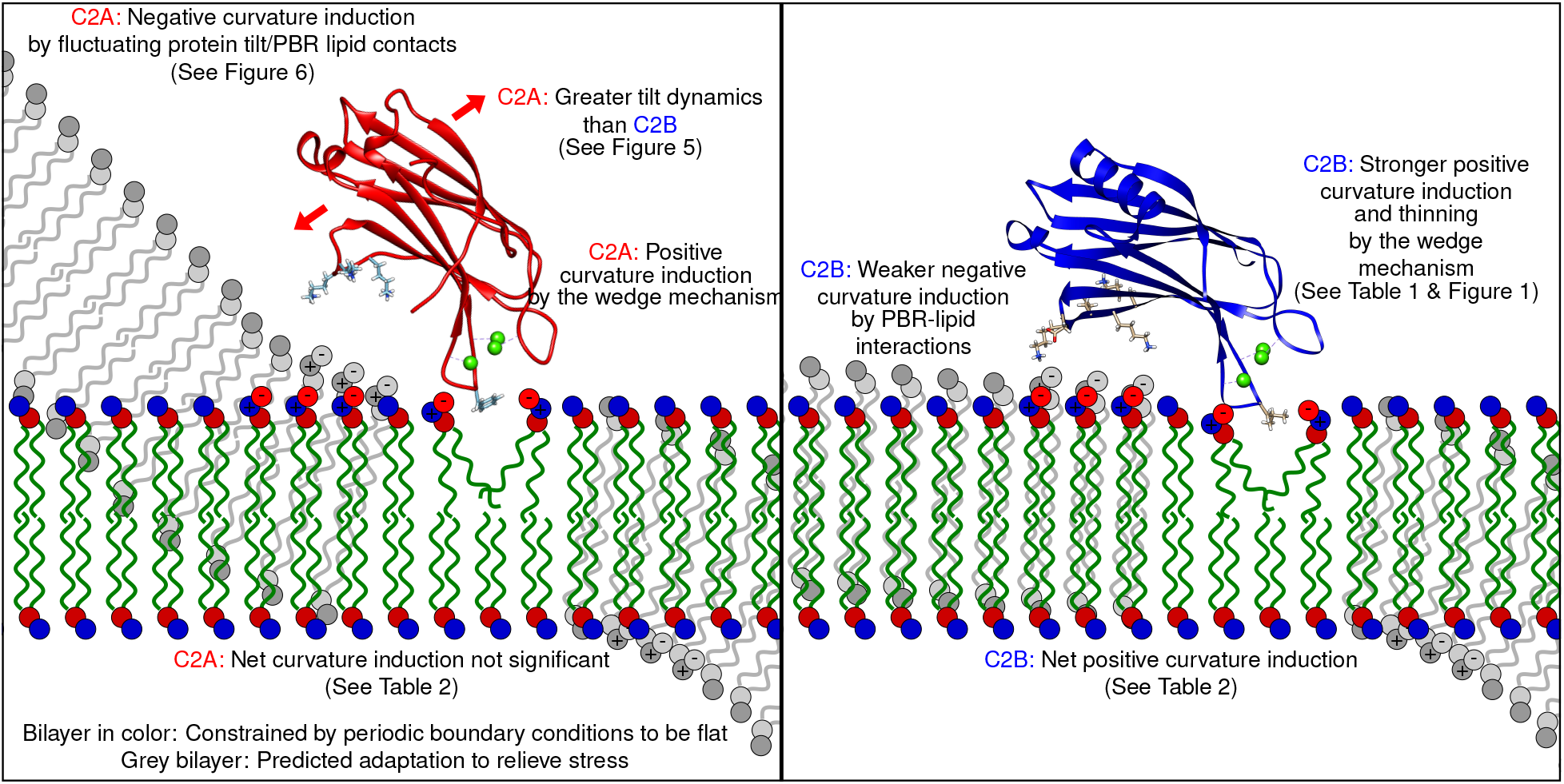
A model of Syt-7 curvature scaffolding in the absence of cholesterol. Inclusion of cholesterol in the membrane may shift C2B more toward the paradigm illustrated on the left.

**Figure 10.**
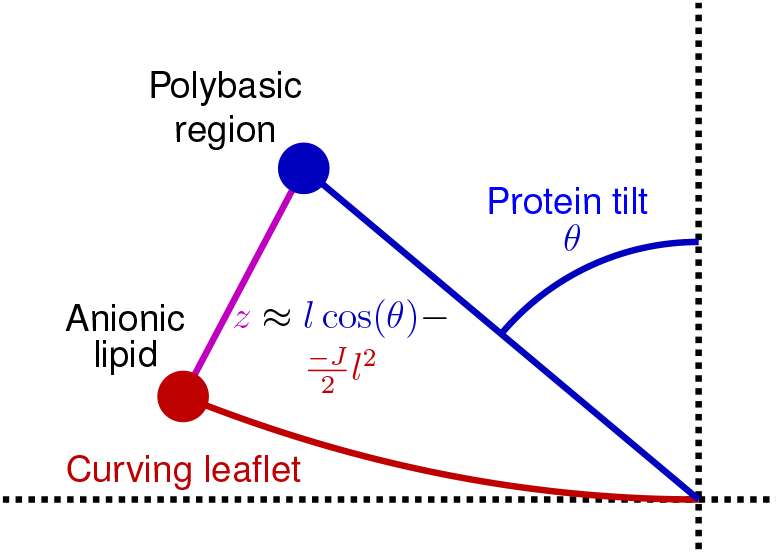
Schematic of the modeled relationship between the PBR/surface distance (*z*), protein tilt (*θ*), and curvature (*J*). Note that by the common sign convention for leaflet curvature, negative curvature is concave with respect to the leaflet (curving up in this schematic), negating *J*. The approximation described is valid when *J* is small and the protein is highly tilted.

#### Curvature induction by the wedge mechanism

Campelo, McMahon and Kozlov (44) estimate the spontaneous curvature of an AAH via the wedge mechanism, validated by subsequent all-atom simulations using the LPP method (46). In the continuum mechanical model, the spontaneous curvature of a wedge inclusion is determined by the area fraction and by the amount of space underneath the wedge that the surround lipids must fill. Both the depth and area are required to estimate the curvature force. Here *thinning* serves as a proxy for the space created beneath the CBL insertion. Consider from **Figure 2** that both C2 domains impact a ∼10 Å radius area at the insertion point, and that the thinning by C2B is approximately 2 Å greater (**Table 1**). From Figure 7a of Campelo et al. (44), a 2 Å difference in thinning effect may lead to as much as a 0.04 Å^−1^ difference in spontaneous curvature (Δ*J*_0,wedge_). Considering the area fraction 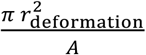 (with simulation area 9000 Å^2^ the deformation area is 7% of the simulation box), the difference of impact on 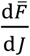 between C2B and C2A due to the wedge mechanism,

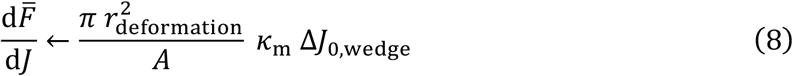

is ∼0.02 kcal/mol/Å, assuming the leaflet bending modulus *k*_m_= 7 kcal/mol.

The wedge and tilt-coupling mechanisms are thus predicted to be of similar magnitude. Although it is difficult to decouple the two effects, we see that C2B has relatively little tilt-coupling with its primary PBR (see **Figure 8**), and thus we attribute its *positive* curvature induction effect to the wedge mechanism. In the case of C2A, we attribute its negligible effect on *total* curvature induction to be a result of the cancellation of tilt-coupling by its modest wedging effect.

## 5. CONCLUSIONS

The simulated structure and fluctuations of synaptotagmin-7 (Syt-7) C2 domains (C2A, C2B, and tandem C2AB) bound to anionic lipid membranes are combined with curvature free energy derivatives from lateral pressure profiles (LPPs; **Table 2**) to yield models of Syt-7’s curvature-generating force. Two mechanisms are proposed (**Figure 9**). First, lysine and arginine residues on the flanks of the C2 domains (poly-basic regions; PBRs) dynamically bind the anionic surface to the protein, inducing negative curvature. Second, the leaflet-embedded calcium binding loops (CBLs) induce positive curvature by a wedging mechanism similar to amphipathic alpha-helices. While both mechanisms are at play for the C2A and C2B domains, tilting and PBR contacts were strongly correlated for C2A (**Figure 8**), counteracting the wedge mechanism when averaged over the simulated bilayer patch, as quantified by LPPs. Anionic lipids enrich at the PBR and CBL binding sites (**Figures 2 and 3**), providing the binding affinity driving both mechanisms. The modeled curvature induction is consistent with experimental results on Syt-1 (84), however, the differences in membrane affinity make direct comparisons impossible. For C2B, a cholesterol-rich bilayer reduced wedging, suggesting that membrane cholesterol can fill the void under C2B and relieve lipid tail stress. This mechanism is consistent with a FRET-based assay of C2 domain bilayer-binding that indicate cholesterol strengthens the C2B-bilayer interaction (**Figure 4**).

The model and structural observations provide a quantitative guide for determining the impact of a Syt protein on fusion pore stability. The fusion pore neck has anisotropic saddle curvature – positive curvature in one direction and negative in the other. When measured at the leaflet surface, the fusion pore has total negative curvature in *both* leaflets as well as a thin hydrophobic interior (53). The range of curvature induction predicted by the tandem Syt C2 domains (spatially distinct positive and negative induction sites) as well as thinning by the CBL wedge, indicate that Syts should support highly curved, anisotropic fusion pore structures. Furthermore, the Syt-7 C2B domain has an arginine apex expected to induce negative curvature on fusion pores (its interactions could not be sampled in this work’s planar simulations). The simulated C2B domain embeds even deeper in tandem (**Table 1**) suggesting the connection between the domains (see **Figure S13** for connection angle distributions) may be important for pore stabilization. Simulating Syt-7 proteins on fusion pores will further validate the structural predictions of this work – allowing the observation of localization of the CBLs to thin regions of the neck, the relaxation of both tilt and PBR contacts, and the arginine apex contact with the membrane.

## 6. ACKNOWLEDGMENTS

This work was supported by the Intramural Research Program (IRP) of the *Eunice Kennedy Shriver* National Institute of Child Health and Human Development (NICHD) at the National Institutes of Health (NIH). AHB was supported by a Postdoctoral Research Associate Training (PRAT) fellowship from the National Institutes of General Medical Sciences (NIGMS), award number 1Fi2GM137844-01. Additional support came from the Camille and Henry Dreyfus Foundation, award TH-18-061 to JDK, and from a CU Denver Eureca Fellowship to VB. This work utilized computational resources of the NIH HPC Biowulf cluster (http://hpc.nih.gov) and resources provided by the NICHD IRP.

## Notes

### Competing Interest Statement

The authors have declared no competing interest.

