## Supporting Material for "Synaptotagmin 7 C2 domains induce membrane curvature stress via electrostatic interactions and the wedge mechanism"

### A. Systems built and simulated.

| System | Lipid-only | C2A | C2B | C2AB |
|---|---|---|---|---|
| PC:PS | Y | Y | Y | Y |
| PC:$PIP_2$ | Y | Y | Y | Y |
| PC:PS:$PIP_2$ | Y | Y | Y | Y |
| PC:PS:cholesterol | Y | Y | Y | Y |

**Table 1.** Planar bilayer systems simulated using the CHARMM all-atom force field. Lipid-only PC:PS, PC:PIP2, and PC:PS were single replica simulations that were run for 400 ns apiece with minimal pre-equilibration using NAMD and a 1 fs timestep. PC:PS:cholesterol was simulated in triplicate for 165 ns apiece with minimal pre-equilibration using NAMD and a 1 fs timestep. For systems including protein, all were simulated in triplicate and had 2 µs equilibration time (using Amber and a 2 fs timestep). All protein-bilayer systems were simulated a further 200 ns (using NAMD and a 1 fs timestep) for dFdR calculation.

### B. Initial setup and orientation of Syt-7 C2 domains on a planar bilayer.

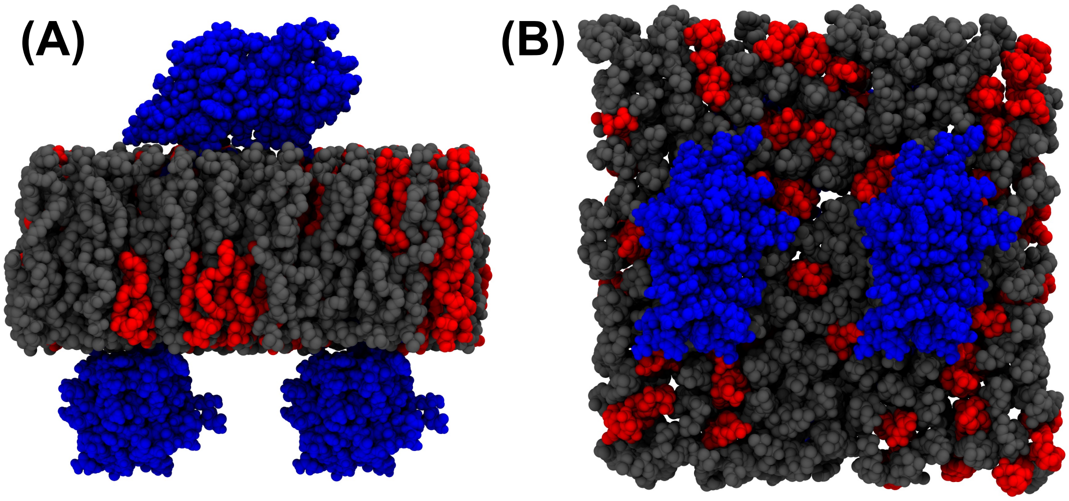


**Figure S2. (A)** Side and **(B)** top views of the initial conditions. This example of C2A (blue) on a PC:PS bilayer (PC is grey and PS is red, respectively).

### C. Line model of POPC lipid.

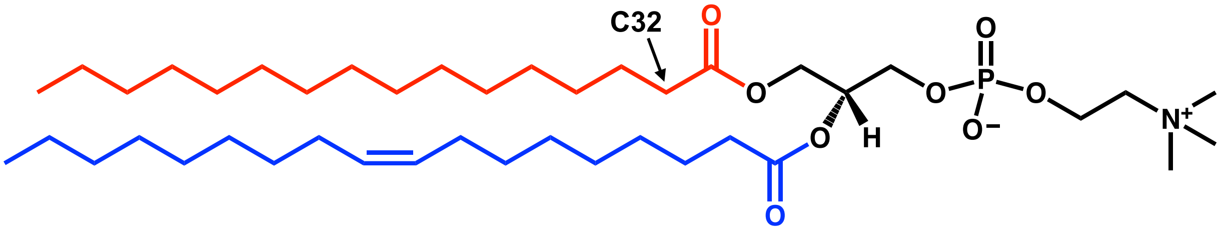


**Figure S2.** The location of the C32 atom in the *sn-2* chain (red). The *sn-1* chain is colored blue.

### D. 2D plots.

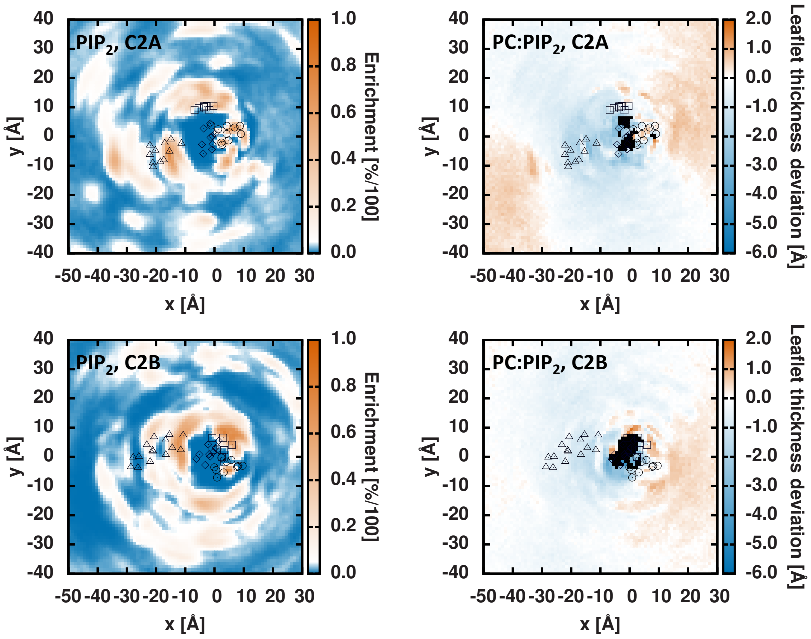


**Figure S3.** C2A (top) and C2B (bottom) in PC:$PIP_2$ membranes. (Left) $PIP_2$ enrichment relative to the bulk value of 5 mol%. (Right) Leaflet thickness relative to the bulk, lipid-only thickness. For C2A: circles are residues 163–172, triangles are residues 176–186, squares are residues 192–198, and diamonds are residues 225–233. For C2B: circles are residues 293–303, triangles are residues 307–321, squares are residues 325–331, and diamonds are residues 357–365. Black pixels represent bins with total density < 20% of the bulk density.

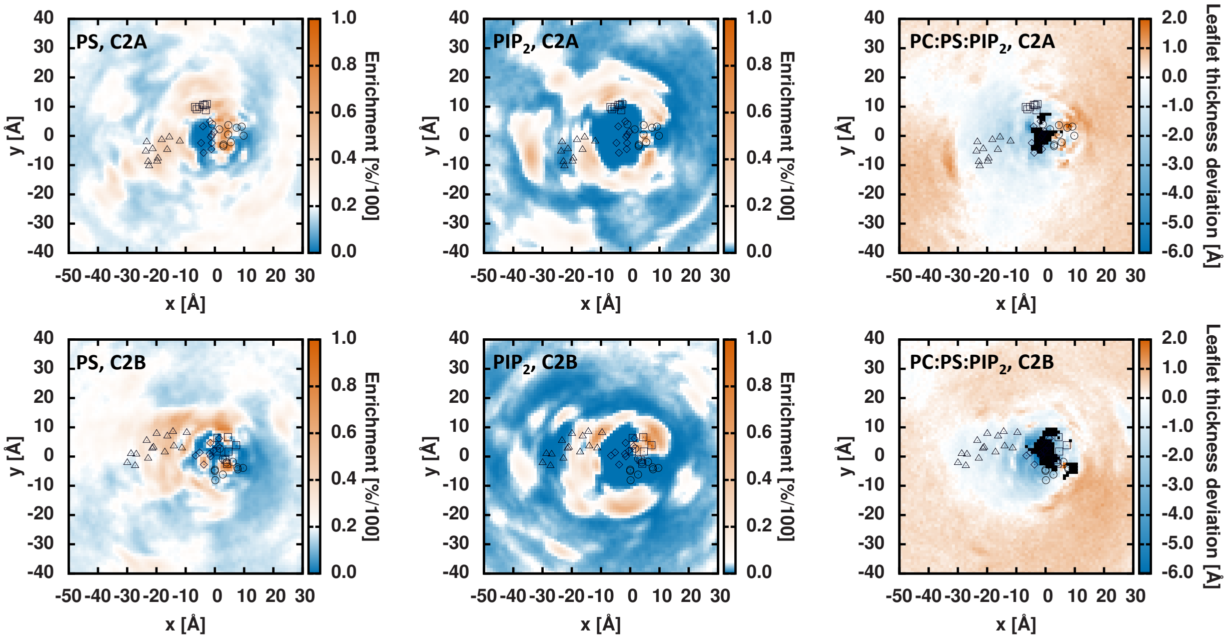


**Figure S4.** C2A (top) and C2B (bottom) in PC:PS:$PIP_2$ membranes. (Left) PS enrichment relative to the bulk value of ~20 mol%. (Middle left) $PIP_2$ enrichment relative to the bulk value of ~5 mol%. (Right) Leaflet thickness relative to the bulk, lipid-only thickness. For C2A: circles are residues 163–172, triangles are residues 176–186, squares are residues 192–198, and diamonds are residues 225–233. For C2B: circles are residues 293–303, triangles are residues 307–321, squares are residues 325–331, and diamonds are residues 357–365. Black pixels represent bins with total density < 20% of the bulk density.

### E. Lipid density along *y* = 0 and lipid tail splay.

(Left) Cross-sections through the total leaflet lipid density. The cross-section is along the *y* = 0 Å axis, averaging the *y* = 0 Å bin and surrounding bins to create a 5 Å cross-section. This was done to reduce noise. (Right) Semi-quantitative lipid tilt caused by the C2 domains. The head and tail of each vector reflects the head and tail of lipids in the bin. The length of the vector reflects the degree of tilt. The vector direction reflects the average tilt direction of lipids in the bin.

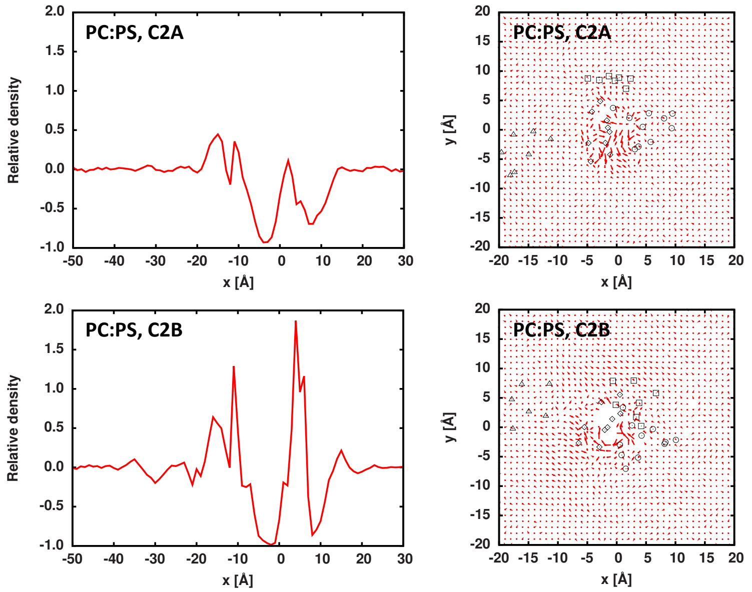


**Figure S5.** (Top) C2A and (bottom) C2B in PC;PS membranes. (Left) A cross-section of the total lipid density along the *y* = 0 Å axis. (Right) Semi-quantitative lipid tilt caused by the C2 domains.

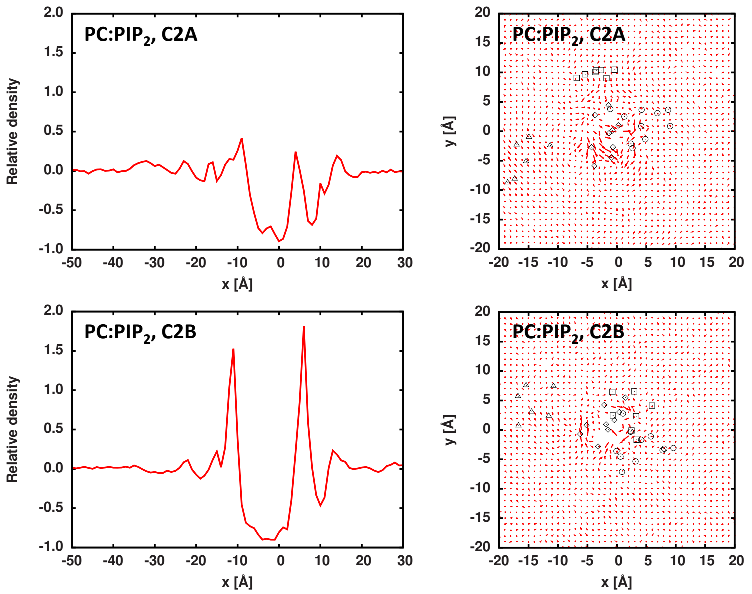


**Figure S6.** (Top) C2A and (bottom) C2B in PC:$PIP_2$ membranes. (Left) A cross-section of the total lipid density along the *y* = 0 Å axis. (Right) Semi-quantitative lipid tilt caused by the C2 domains.

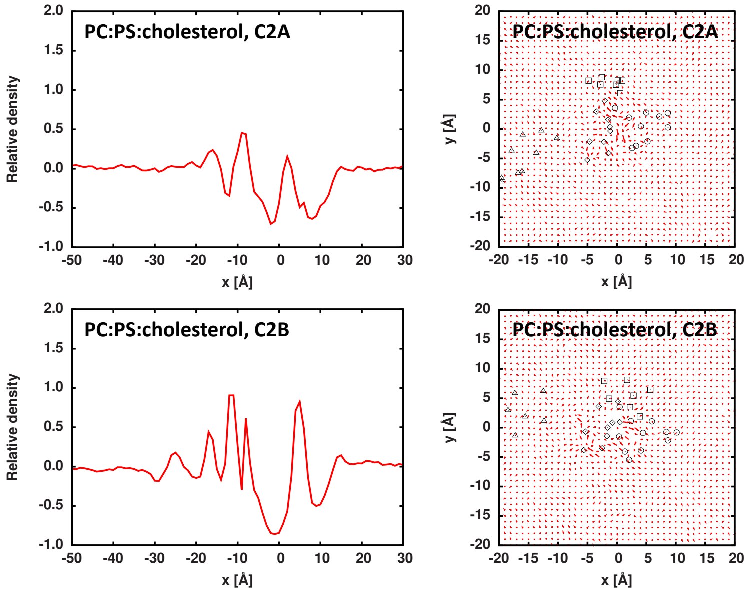


**Figure S7.** (Top) C2A and (bottom) C2B in PC:PS:chol membranes. (Left) A cross-section of the total lipid density along the $y$ = 0 Å axis. (Right) Semi-quantitative lipid tilt caused by the C2 domains.

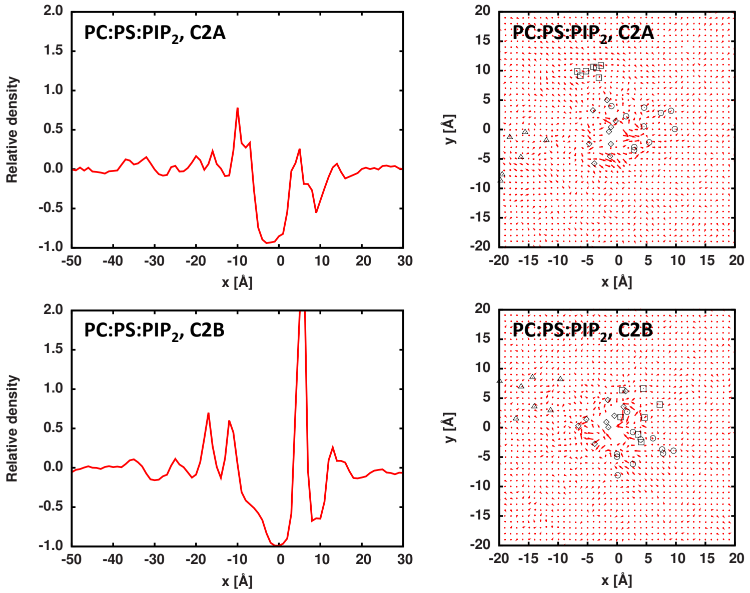


**Figure S8.** (Top) C2A and (bottom) C2B in PC:PS:$PIP_2$ membranes. (Left) A cross-section of the total lipid density along the $y$ = 0 Å axis. (Right) Semi-quantitative lipid tilt caused by the C2 domains.

### F. Per-residue contacts with the environment.

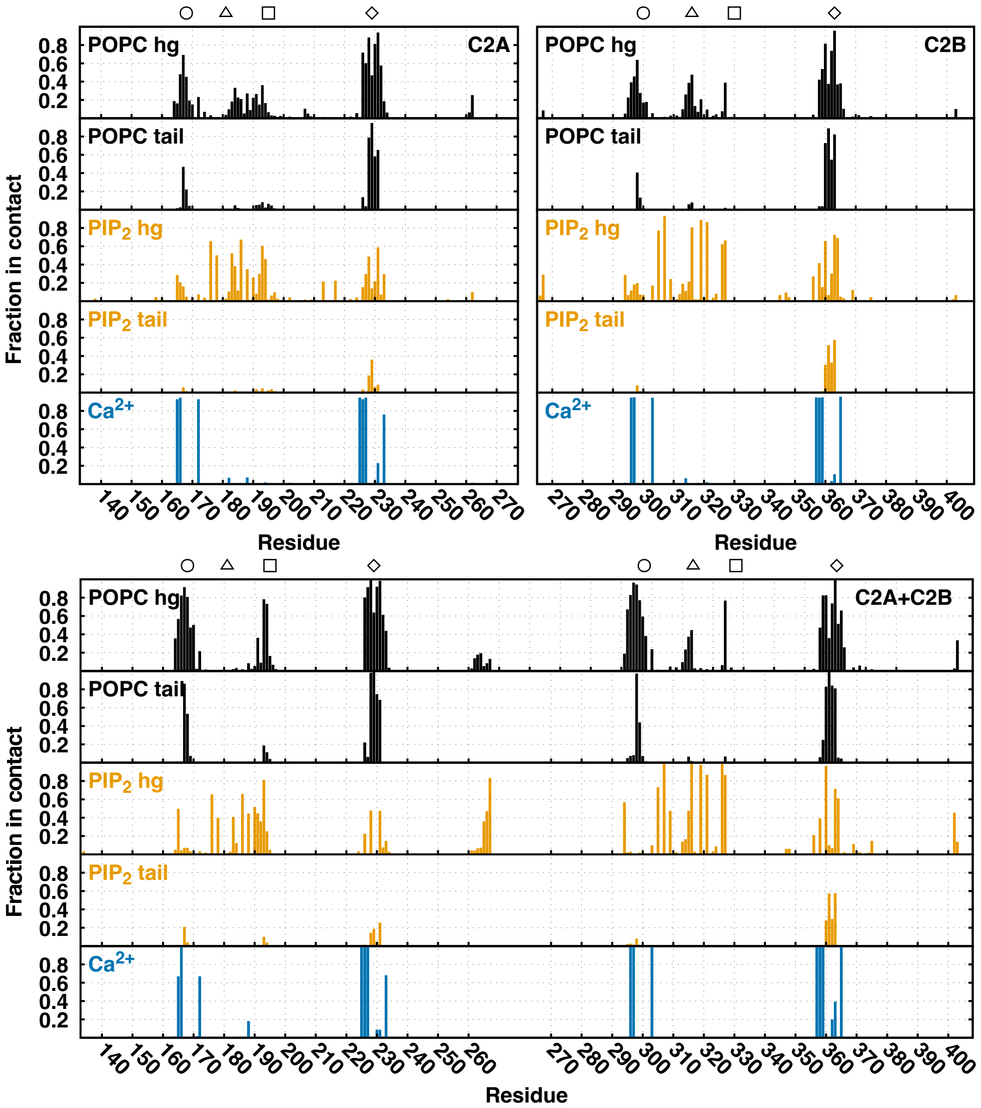


**Figure S9.** PC:$PIP_2$ contact plots for solo C2A (left), solo C2B (middle), and tandem C2A+C2B (right). PC head group (hg) and tail contacts are in black, PIP2 head group and tail contacts are in orange, and calcium contacts are in blue. The circle, triangle, square, and diamond positions correspond to residues 168, 181, 195, and 229 for C2A and residues 298, 314, 328, and 361 for C2B. A value of "0" indicates an unobserved interaction through our entire ensemble, and a value of "1" indicates that the interaction was present 100% of the time in our entire ensemble.

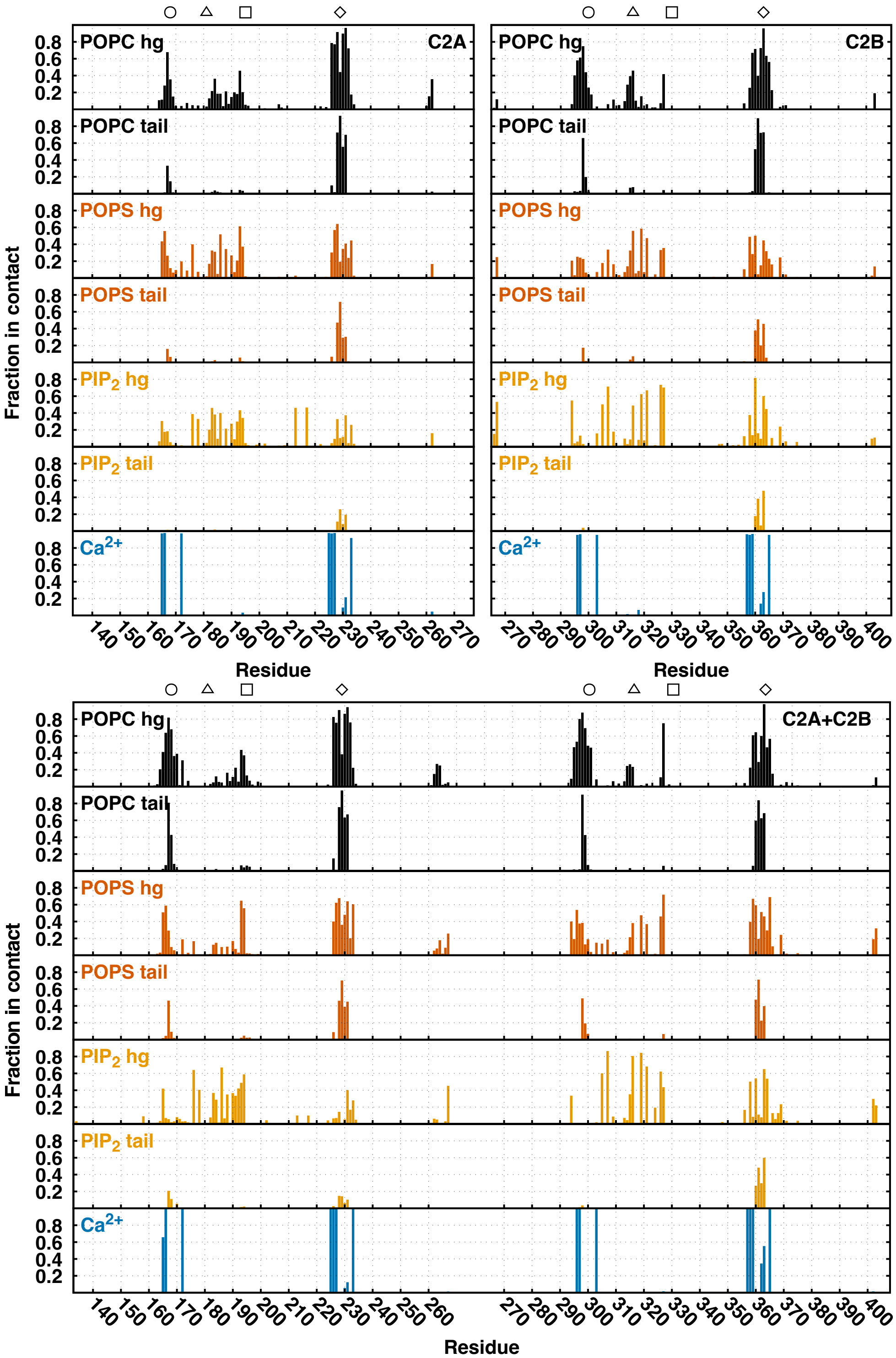


**Figure S10.** PC:PS:$PIP_2$ contact plots for solo C2A (left), solo C2B (middle), and tandem C2A+C2B (right). PC head group (hg) and tail contacts are in black, PS head group and tail contacts are in vermillion, $PIP_2$ head group and tail contacts are in orange, and calcium contacts are in blue. The circle, triangle, square, and diamond positions correspond to residues 168, 181, 195, and 229 for C2A and residues 298, 314, 328, and 361 for C2B. A value of "0" indicates an unobserved interaction through our entire ensemble, and a value of "1" indicates that the interaction was present 100% of the time in our entire ensemble.

### G. Membrane contacts.

The residue contacts with lipid head groups from **Figures 5–6** were cumulatively summed to get the total number of protein-lipid contacts. The cumulative standard error from the three replicas is the square root of the summed error.

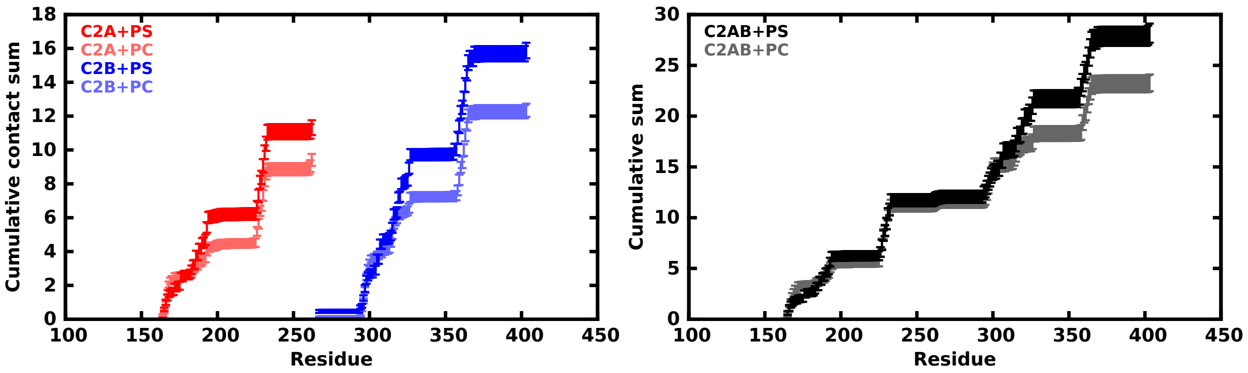


**Figure S11.** C2 domain summed contacts with PC:PS lipid head groups. As a solo C2A domain, Syt-7 forms 11.3 ± 1.6 contacts with PS and 9.4 ± 1.3 contacts with PC. As a solo C2B domain, Syt-7 forms 15.9 ± 1.6 contacts with PS and 12.3 ± 1.5 contacts with PC. As a C2AB tandem, Syt-7 forms 28.2 ± 2.7 contacts with PS and 23.3 ± 2.4 contacts with PC.

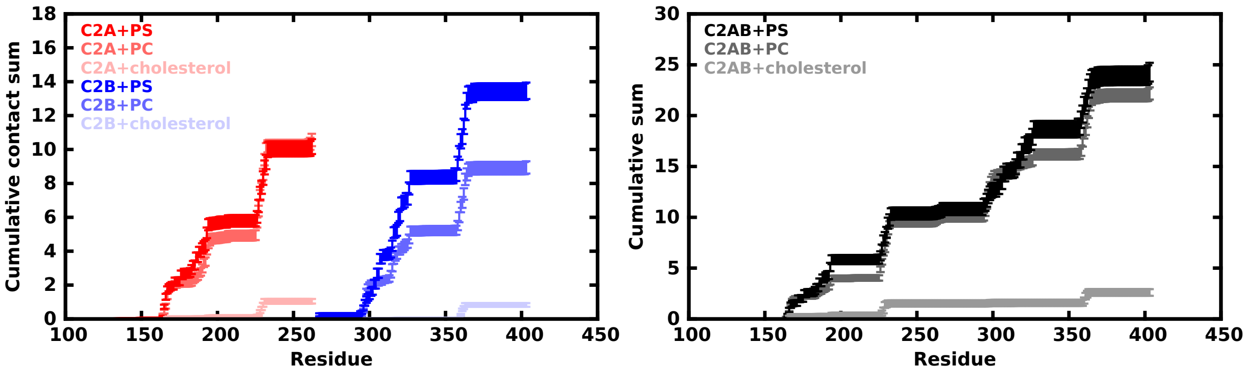


**Figure S12.** C2 domain summed contacts with PC:PS:cholesterol lipid head groups. As a solo C2A domain, Syt-7 forms 10.2 ± 1.6 contacts with PS, 10.6 ± 1.5 contacts with PC, and 0.9 ± 0.5 contacts with cholesterol. As a solo C2B domain, Syt-7 forms 13.5 ± 1.7 contacts with PS, 8.9 ± 1.4 contacts with PC, and 0.8 ± 0.5 contacts with cholesterol. As a C2AB tandem, Syt-7 forms 24.3 ± 2.6 contacts with PS, 22.2 ± 2.1 contacts with PC, and 2.6 ± 1.0 contacts with cholesterol.

**H. C2AB tandem orientation projected on the *xy*-plane.**

The C2AB tandem is composed of C2A and C2B domains connected by a short, but flexible linker that allows dynamic reorientation of the C2 domains. To define the C2AB tandem orientation, individual C2A and C2B vectors were defined from the $CoM_{CBL}$ to the $CoM_{C2}$ projected onto the *xy*-plane. The angle between these vectors was then determined by $\text{atan2}(-\text{det}, -\text{dot}) + \pi$, where dot and det are the dot product and determinant of the vectors, respectively (range of $[0, 2\pi]$). This angle becomes poorly defined when a domain is perpendicular to the membrane surface, but these orientations are rare according to **Figure 1**. The angles were calculated over the last 1 μs of trajectory for all replicas.

Connection angles centered near 180° indicate that domains typically orient in an anti-parallel ("extended") conformation. Forays toward smaller/larger angles indicate parallel ("side-by-side") orientations. Adding cholesterol significantly shifts the connection angle distribution to explore more parallel conformations. Three of the six C2AB total tandems (i.e., two C2AB tandems apiece in three replicas) visited < 50° conformations in the PC:PS:cholesterol membrane. Each C2AB tandem that samples the < 50° conformation exits at least once. The C2AB tandem in PC:PS:$PIP_2$ composition explores wider angles than the other compositions. In the case of the PC:PS:cholesterol membrane, we hypothesize that the reduced deformation of Syt-7 in cholesterol-containing membranes shifts the C2AB tandem orientation.

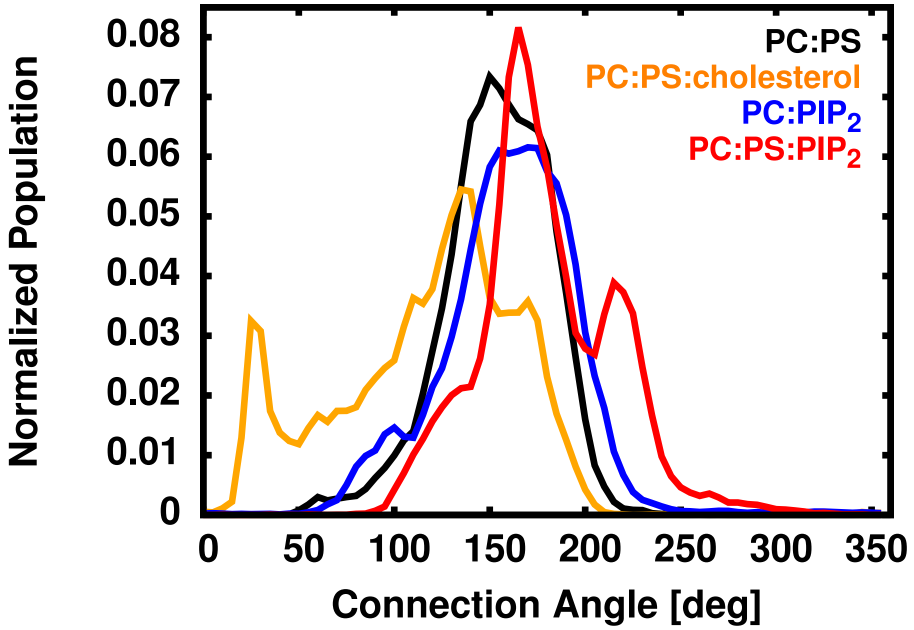


**Figure S13.** Normalized connection angle between the C2 domain components of the C2AB tandem.
